# Gener*anno*: A Genomic Foundation Model for Metagenomic Annotation

**DOI:** 10.1101/2025.06.04.656517

**Authors:** Qiuyi Li, Wei Wu, Yiheng Zhu, Fuli Feng, Jieping Ye, Zheng Wang

**Affiliations:** Alibaba Cloud Computing, Beijing, China; Zhejiang University, Hangzhou, China; Institute of Dataspace, Hefei, China

**Keywords:** Genomic Foundation Model, Metagenomics, Gene Annotation, Taxonomic Classification

## Abstract

The rapid growth of genomic and metagenomic data has underscored the pressing need for advanced computational tools capable of deciphering complex biological sequences. In this study, we introduce **Gener***anno*, a compact yet powerful genomic foundation model (GFM) specifically optimized for metagenomic annotation. Trained on an extensive dataset comprising 715 billion base pairs (bp) of prokaryotic DNA, **Gener***anno* employs a transformer encoder architecture with 500 million parameters, enabling bidirectional attention over sequences up to 8192 bp at single-nucleotide resolution. This design addresses key limitations of existing methods, including the inability of traditional Hidden Markov Models (HMMs) to handle fragmented DNA sequences from multi-species microbial communities, as well as the suboptimal tokenization schemes of existing GFMs that compromise fine-grained analysis. At its core, **Gener***anno* excels in identifying coding regions from fragmented and mixed DNA sequences—a hallmark of metagenomic analysis. It achieves superior accuracy compared to traditional HMM-based methods (e.g., GLIMMER3, GeneMarkS2, Prodigal) and recent LLM-based approaches (e.g., GeneLM), while demonstrating robust generalization ability on archaeal genomes. Leveraging its advanced contextual understanding capability, **Gener***anno* further enables two essential functions: pseudogene prediction and taxonomic classification—both performed based solely on raw sequence data, without reliance on reference databases or comparative genomics. These functionalities collectively streamline the metagenomic analysis pipeline, significantly reducing preprocessing requirements and enabling end-to-end interpretation of sequencing data. Beyond its primary role in metagenomic annotation, **Gener***anno* also serves as a powerful GFM. To evaluate its broader utility, we curated the Prokaryotic Gener Tasks—a comprehensive benchmark suite specifically tailored for prokaryotic genomic analysis. It includes gene fitness prediction, antibiotic resistance identification, gene classification, and taxonomic classification, reflecting diverse aspects of functional genomics. On this benchmark, **Gener***anno* consistently outperforms existing GFMs such as DNABERT-2, NT-v2, and GenomeOcean, demonstrating strong generalization capabilities across a wide range of genomic tasks. Overall, **Gener***anno* provides a unified framework that integrates multiple critical functions for metagenomic annotation and beyond. By eliminating dependencies on external resources and offering rich contextual understanding of genomic sequences, this work delivers a foundational tool for advancing functional genomics in complex microbial communities. Implementation details and supplementary resources are available at https://github.com/GenerTeam/GENERanno.

## 1 Introduction

In recent years, the rapid advancement of high-throughput sequencing technologies [61] has enabled the collection of genetic data from diverse organisms on an unprecedented scale. This wealth of genomic information has significantly enhanced our understanding of biological systems, providing crucial insights into genome structure, function, and evolution. These developments have profound implications for human health, agriculture, and environmental sustainability. However, the vast scale and complexity of these datasets present significant computational challenges.

Metagenomics [37], in particular, has emerged as a cornerstone of modern biology, offering invaluable insights into the collective genetic material of microbial communities. These communities are vital to ecosystems, human health, and industrial applications. Traditional methods for prokaryotic gene prediction and annotation, such as GLIMMER [25, 26, 27], GeneMark [9, 10, 47], and Prodigal [38], have long been recognized as the gold standard in prokaryotic genomic analysis. These approaches, typically based on Hidden Markov Models (HMMs) [59], are adept at identifying coding regions in assembled prokaryotic genomes. However, they frequently encounter difficulties in metagenomic settings, where sequences are not only short and fragmented but also originate from highly complex, multi-species microbial communities. Analyzing such fragmented and mixed DNA sequences remains a formidable challenge due to their inherent complexity, diversity, and sheer abundance. This difficulty arises from the reliance on rigid statistical models that lack the adaptability to interpret the nuanced contextual complexities of metagenomic data, underscoring the pressing need for more versatile tools capable of directly analyzing fragmented DNA while effectively capturing intricate biological patterns across diverse genomic contexts.

In response to these challenges, recent advancements in machine learning, particularly the development of large language models (LLMs) [80], have gained traction in genomics research. There has been an extensive literature of leveraging LLMs for analysing biological sequences [1, 35, 20, 13]. These models employ large-scale unsupervised pre-training to capture intricate biological patterns and generalize across diverse tasks. In genomics, they are often referred to as genomic foundation models (GFMs). There are two broad classes of LLMs utilized for genomic research. Masked language models (MLM) [28] feature bidirectional attention and are well-recognized for their sequence understanding abilities. Examples include DNABERT [40, 82], Nucleotide Transformer (NT) [20], LucaOne [36], GROVER [64], Caduceus [66], and GENA-LM [30]. While early efforts in developing genomic language models primarily focused on MLM, causal language models (CLM) [2] have become increasingly popular recently. CLM-based models feature causal attention, focusing only on preceding contexts, and are renowned for their generative capabilities while preserving sequence understanding. Models like HyenaDNA [52], megaDNA [67], Evo [51, 13], METAGENE [46], GenomeOcean [83], **Gener***ator* [78], and HybriDNA [49] fall into this category. Notably, another emerging class of generative GFMs is based on diffusion models [19], such as D3 [65], MDLM [62], and DDSM [6]. These models operate by iteratively denoising data, starting from random noise and progressively refining it into structured outputs. This approach offers robustness in complex sequence design tasks. However, in this study, we focus primarily on CLM-based and MLM-based GFMs due to their widespread adoption and established performance in genomic tasks.

We advocate for a reconsideration of the growing preference for CLM over MLM in genomic modeling. Both approaches possess distinct strengths suited for different applications. CLM models excel in DNA sequence design and optimization, leveraging their generative capabilities to enable precise genomic interventions [51, 78]. In contrast, MLM models are particularly well-suited for tasks requiring bidirectional contextual information, such as gene annotation [70]. Gene annotation, forming the foundation of genomic and proteomic analysis, aims to accurately identify protein-coding regions within DNA sequences. This task differs significantly from general sequence understanding, as reflected in established genomic benchmarks. To clarify this distinction, general sequence understanding typically involves coarse-grained analyses of entire sequences. Examples include tasks such as sequence classification (e.g., gene classification [78]) and sequence regression (e.g., enhancer activity prediction [23]), where both MLM and CLM models perform comparably well. By contrast, gene annotation requires fine-grained nucleotide-level analysis—specifically, determining whether each nucleotide belongs to a coding region.

Several attempts have been made to leverage MLM-based genomic foundation models for gene annotation, with notable examples including SegmentNT [24] and GeneLM [3]. SegmentNT is an annotation method built upon the Nucleotide Transformer (NT). Specifically, SegmentNT-Human is tailored for annotating the human genome, while SegmentNT-Multispecies is trained on five selected animal species (mouse, chicken, fly, zebrafish, and worm) and demonstrates a degree of generalizability across other eukaryotic genomes. GeneLM, akin to **Gener***anno*, targets prokaryotic metagenomic annotation. Built upon DNABERT, GeneLM is trained on a comprehensive collection of prokaryotic reference genomes and demonstrates relatively robust performance across various prokaryotic genomic annotation tasks. Despite their promise, these methods face substantial limitations that impede their practical utility. Notably, they often struggle to consistently outperform traditional HMM-based approaches, despite the inherent ability of LLMs to capture intricate biological patterns. Additionally, the computational demands of these approaches further restrict their scalability in large-scale applications. Consequently, while these methods represent promising initial efforts, they fall short of delivering the transformative improvements expected from LLM-based genomic tools. These limitations primarily stem from two aspects: first, the inherent performance constraints of the underlying genomic foundation models; and second, the suboptimal configuration of these models for gene annotation tasks. Specifically, **Gener***anno*: A Genomic Foundation Model for Metagenomic Annotation the application of tokenizers in genomic foundation models inadvertently compromises single-nucleotide resolution, which is crucial for gene annotation tasks.

Given this context, we introduce **Gener***anno*, a genomic foundation model specifically optimized for gene annotation tasks. This model employs a transformer encoder architecture with 500 million parameters and a context length of up to 8192 base pairs (bp) at single-nucleotide resolution. Trained on an expansive dataset comprising 715 billion base pairs of prokaryotic DNA, the extensive and diverse pre-training data endow **Gener***anno* with enhanced capabilities for understanding genomic contexts across a wide array of organisms. A noteworthy aspect of our research is the recognition of the absence of standardized benchmark metrics in the prokaryotic domain, akin to established ones in the eukaryotic domain such as the NT tasks [20], GUE [82], Genomic Benchmark [32] and BEND [50]. In response, we curated and assembled a collection of biologically meaningful tasks in the prokaryotic domain, termed the Prokaryotic Gener Tasks. Addressing a critical gap in the field, the Prokaryotic Gener Tasks comprise four key components: gene fitness prediction under various experimental conditions, antibiotic resistance prediction, gene classification, and taxonomic classification. Through comprehensive benchmark evaluations, **Gener***anno* consistently surpasses its counterparts, such as NT-v2, DNABERT-2, and GenomeOcean. This superiority firmly establishes **Gener***anno* as an outstanding genomic foundation model in the prokaryotic domain, laying the groundwork for its exceptional performance in metagenomic annotation.

For metagenomic annotation [63], we tested the model performance on a comprehensive dataset comprising 33 distinct prokaryotic species, including both genome and plasmid sequences. In comparative analyses with widely adopted state-of-the-art gene annotation methods such as GLIMMER3, GeneMarkS2, Prodigal, and GeneLM, **Gener***anno* consistently demonstrates substantial superiority, positioning it among the leading methods for both genomic and metagenomic annotation. We further assessed the zero-shot predictive capabilities of **Gener***anno* on archaeal genomes. Remarkably, the model performance matched the ‘in-sample’ predictive accuracy of traditional HMM-based methods, significantly surpassing GeneLM, thereby demonstrating exceptional generalization ability. Beyond standard annotations, **Gener***anno* pioneers the prediction of pseudogenes based solely on sequence data. Pseudogenes are non-functional DNA sequences that resemble functional genes but have lost their ability to code for proteins due to mutations or genomic rearrangements. Traditional methods typically rely on comparative genomics and functional assays to identify pseudogenes [72], which can be time-consuming and require comprehensive reference databases. In contrast, **Gener***anno* offers a streamlined and efficient solution, leveraging advanced contextual understanding to differentiate pseudogenes from active coding sequences.

Overall, **Gener***anno* emerges as a compact yet powerful genomic foundation model within the prokaryotic domain. Across a comprehensive benchmark comprising a suite of biologically meaningful tasks, it consistently demonstrates strong performance, underscoring its adeptness in capturing intricate biological patterns. Beyond its excellence as a genomic foundation model, **Gener***anno* is meticulously optimized for metagenomic annotation tasks, consistently surpassing traditional HMM-based gene annotation methods. This remarkable performance highlights the potential of LLMs in revolutionizing gene annotation practices. **Gener***anno* not only sets a new standard in genomic analysis but also paves the way for the next generation of gene annotation methodologies. This noteworthy achievement underscores the transformative potential of large language models in evolving gene annotation practices. By setting a new benchmark in genomic analysis, **Gener***anno* contributes to laying the groundwork for the future development of advanced gene annotation technologies.

## 2 Method

### 2.1 Data Preparation

For model pre-training, we sourced raw DNA sequences from all prokaryotic organisms in the RefSeq database [54]. In our prior work on **Gener***ator* [78], we introduced and validated a data curation strategy termed *Functional Sequence Training*. Our experimental results demonstrated that this approach significantly enhances the performance of pre-trained models compared to indiscriminately using all genomic sequences for training. This finding was further corroborated by Evo2 [13], which reported similar conclusions. Therefore, in this study, we continued to adopt the functional sequence training strategy for **Gener***anno*. Specifically, leveraging the extensive annotation data available in RefSeq, we extracted biologically functional regions from genomic sequences. These regions encompass a broad spectrum of functionalities, including transcription into various RNA molecules, translation into complex proteins, and regulatory functions such as promoters and enhancers that govern gene expression. Collectively, these functional DNA segments constituted our training dataset, totaling 715 billion nucleotides.

The rationale for adopting functional sequence training lies in the inherent randomness of genetic mutations, which renders DNA far from being a concise language. At the origin of life, DNA likely began as nearly random sequences, and through stochastic mutations and other evolutionary events [43, 44], functional regions emerged. This process is analogous to randomly typing on a keyboard, where there is a small probability of producing a coherent sentence. Such ‘sentences’ (functional regions) are retained due to their selective advantages and have accumulated over billions of years of evolution, ultimately leading to the extant biodiversity. The stochastic nature of genetic mutations also implies that functional regions are sparse and interspersed within vast stretches of non-functional DNA, often referred to as ‘junk DNA’ [12]. While some of these regions may harbor undiscovered genes, the majority exhibit significantly higher mutation rates compared to functional regions [5], as they are not subject to selective pressures. In fact, experimental evidence shows that even when portions of this non-functional DNA are removed via genome editing, organisms can still maintain normal biological functions [58]. Therefore, including non-functional DNA in the training set does not effectively increase data volume but rather introduces noise, potentially degrading the quality of the training data and, consequently, model performance.

### 2.2 Tokenization

Tokenization is a fundamental step in natural language processing that involves breaking down input sequences into discrete units, or tokens, which serve as the basic building blocks for model training. Most genomic foundation models (GFMs) strive to employ suitable tokenizers, such as K-mer [17] or Byte Pair Encoding (BPE) [42], which group multiple consecutive nucleotides into a single token, thereby extending the context window. However, this approach inadvertently compromises single-nucleotide resolution, which is crucial for tasks like gene annotation. For example, both SegmentNT (NT) and GeneLM (DNABERT) employ a 6-mer tokenizer, necessitating additional mechanisms to decompose tokens and accurately delineate gene boundaries [24, 3]. This decomposition process can lead to accumulated errors, thereby diminishing the capability of GFMs to fully realize their potential.

In light of the specific requirements of gene annotation tasks, we opted for a single-nucleotide tokenizer in **Gener***anno*, treating each nucleotide (A, T, C, G) as an individual token. This choice ensures high-resolution representation of DNA sequences, preserving critical single-nucleotide details necessary for precise gene boundary detection. While some existing GFMs also adopt single-nucleotide tokenizers, they often face limitations in handling long sequences due to constraints on context window size [36], or lack robust support for bidirectional attention [52, 67, 51, 13, 49]. These challenges underscore one of the key motivations behind the development of **Gener***anno*.

It is worth noting that the scalability of transformer models is inherently constrained by the quadratic growth of computational costs associated with attention mechanisms as context length increases. To address this issue, recent studies have explored more streamlined architectures, such as State Space Models (SSMs) [34], exemplified by StripedHyena [57] used in Evo and Mamba [33] in HybriDNA. Despite their efficiency in long-context training, SSMs often struggle to achieve comparable performance in long-context understanding [75, 7]. Additionally, due to their inherent characteristics, SSMs are predominantly designed for causal attention and face challenges in supporting bidirectional attention effectively—a paradigm essential for gene annotation. The BiMamba architecture, as implemented in Caduceus [66], represents a notable exception by combining forward and backward Mamba components. In this study, we retained the transformer encoder architecture for its well-established reliability and compatibility with bidirectional attention. To ensure practical usability, we prioritized maintaining a sufficiently large context window while keeping computational costs within acceptable limits. The detailed model configuration will be provided in the following section.

### 2.3 Pre-training

In terms of model architecture, **Gener***anno* broadly follows the structure of Llama [73], which was originally implemented as a CLM model. We have made self-modifications to support bidirectional attention, aligning it with the MLM paradigm. This adaptation is motivated by the inherent dependence of gene annotation tasks on bidirectional context. For instance, consider the first nucleotide in a sequence: without access to subsequent context, it is impossible to determine whether this nucleotide belongs to a coding region. This highlights the critical role of bidirectional attention in gene annotation, a capability that CLM models inherently lacks.

Notably, our ‘Llama for MLM’ implementation shares similarities with the recently released ModernBERT model [76], both of which are based on the transformer encoder architecture and support the same maximum context length of 8192 tokens. However, ModernBERT employs a hybrid design that combines local and global attention mechanisms in intermediate layers, aiming to improve training efficiency. In contrast, **Gener***anno* exclusively uses global attention across all layers, prioritizing maximal performance for genomic tasks. Although we have not trained or tested **Gener***anno* on natural language processing tasks [76], we reasonably expect that its robust architecture would also yield strong performance in such domains.

The detailed model configuration is provided in Table 1. The pre-training process employs a batch size of 2 million tokens and utilizes the AdamW optimizer [48], coupled with a cosine learning rate scheduler with a warm-up phase. The entire pre-training spans 2 epochs, processing a total of 1.4 trillion tokens. To enhance the efficiency of long-context pre-training, we leverage Flash Attention [22] and the Zero Redundancy Optimizer (ZeRO) [60]. Additional details regarding the pre-training process are provided in the Supplementary Section B. Overall, our configuration effectively harnesses the potential of transformer architectures while being specifically tailored to gene annotation tasks.

**Table 1:**
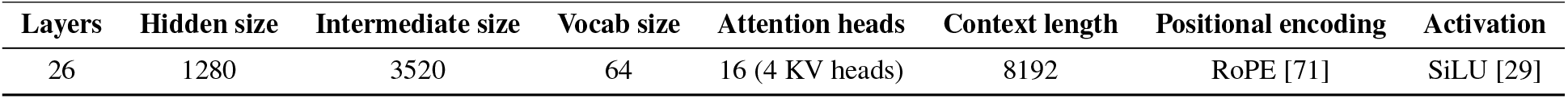
Detailed architecture of Gener*anno*.

| Layers | Hidden size | Intermediate size | Vocab size | Attention heads | Context length | Positional encoding | Activation |
| --- | --- | --- | --- | --- | --- | --- | --- |
| 26 | 1280 | 3520 | 64 | 16 (4 KV heads) | 8192 | RoPE [71] | SiLU [29] |

### 2.4 Prokaryotic Gener Tasks

Beyond the development of **Gener***anno*, another noteworthy contribution of our work is the curation of a biologically meaningful benchmark suite, named Prokaryotic Gener Tasks. While pre-trained genomic foundation models such as Evo and GenomeOcean have demonstrated success in prokaryotic domains, there remains a notable absence of standardized benchmarks akin to NT Tasks [20], GUE [82], Genomic Benchmark [32], or BEND [50]. The establishment of such benchmarks is essential for enabling fair and standardized comparisons between models, thereby fostering advancements in the field.

We conducted a comprehensive analysis of existing eukaryotic genomic benchmarks and identified several limitations. For example, NT Tasks and GUE have been criticized for their limited biological relevance, while Genomic Benchmark primarily focuses on human genomes and suffers from sequence length constraints, prompting the development of Genomic Long-Range Benchmark [74]. Although BEND emphasizes biologically meaningful tasks, it remains restricted to human genomes. These limitations highlight critical challenges in benchmark design. In light of these observations, we aimed to avoid similar pitfalls when curating Prokaryotic Gener Tasks. Specifically, we designed Prokaryotic Gener Tasks with three key principles in mind:

1. **Biological Relevance**: Each task addresses a biologically meaningful question, ensuring that the evaluation reflects real-world applications.
2. **Multispecies Coverage**: The tasks assess both model performance and robustness, ensuring reliable analysis across a wide range of prokaryotic species.
3. **Sequence Length Diversity**: The tasks encompass various sequence lengths to comprehensively evaluate model capabilities across different scales.

Building on these principles, we propose Prokaryotic Gener Tasks, which consist of the following subtasks:

#### Gene Fitness Prediction

This task involves predicting gene fitness scores based on data sourced from the Fitness Browser [58]. Gene fitness measures the importance of a gene for survival under specific experimental conditions. Our benchmark covers diverse environments, including variations in pH levels, temperature, carbon sources, nitrogen sources, and exposure to different chemical compounds. This task is particularly valuable for understanding how genes contribute to microbial adaptability and survival in dynamic environments, providing insights into evolutionary pressures and metabolic pathways.

#### Antibiotic Resistance Prediction

The goal of this task is to predict whether a given gene confers antibiotic resistance. Resistance genes were sourced from the Comprehensive Antibiotic Resistance Database (CARD) [4], while control genes were randomly sampled from RefSeq [54]. To eliminate potential confounding factors, we further adjusted the sampled control sequences to ensure that their length distribution closely matched that of the resistance genes. This task is critical for combating the global challenge of antibiotic resistance by enabling the rapid identification of resistance genes in genomic data.

#### Gene Classification

This task is centered on the classification of genes into distinct functional categories, including coding sequences (CDS), pseudogenes, transfer RNA (tRNA), ribosomal RNA (rRNA), non-coding RNA (ncRNA), and intergenic regions that are either non-functional or of unknown function. The dataset used for this task was carefully balanced and randomly sampled from RefSeq annotations. A key challenge in this task lies in distinguishing between CDS and pseudogenes, as pseudogenes closely resemble functional coding regions but have lost their original functionality due to subtle disruptive mutations or rearrangements. Traditional methods for pseudogene identification often rely heavily on comparative genomics and functional assays [72], which are computationally expensive and require extensive reference databases. In contrast, this task evaluates the capacity of GFMs to predict gene types—including pseudogenes—directly from sequence data. By leveraging their advanced contextual understanding of genomic sequences, GFMs offer a promising alternative to traditional methods, addressing their limitations and providing a more efficient and scalable solution for gene classification.

#### Taxonomic Classification

Taxonomic classification involves identifying organisms based on their genomic sequences [68], a cornerstone of genomic research. Our benchmark utilizes data from the Genome Taxonomy Database (GTDB) [55], which provides comprehensive and standardized taxonomic annotations. In this task, we focus on predicting intermediate taxonomic levels, specifically from phylum to family, while excluding higher (domain) and lower (genus and species) ranks for practical reasons. This decision is motivated by the fact that the domain level (Bacteria) is universal across all samples and thus redundant for prediction, while genus and species levels exhibit fine-grained distinctions but suffer from limited sample sizes, making accurate predictions impractical. By focusing on intermediate levels, we strike a balance between biological relevance and model feasibility, ensuring meaningful and achievable predictions. This task is divided into three subtasks:

1. **SSU Classification**: This subtask focuses on universal marker genes, specifically SSU (e.g., 16S rRNA [8]), which are widely used in taxonomic classification due to their conserved nature and universal presence across species.
2. **Mixed Marker Classification**: In this subtask, one marker gene is randomly selected for each species [56], and predictions are made based on mixed inputs from these marker genes.
3. **Random Fragment Classification**: This subtask involves making predictions directly from randomly sampled genome fragments. Due to the inherent complexity of fragmented genomic sequences, this task is particularly challenging. However, it also holds significant potential for streamlining traditional metagenomic workflows, which typically require assembling fragmented sequences into contigs and identifying marker genes prior to classification.

By emphasizing biological relevance, multispecies coverage, and sequence length diversity, we aim to establish a standardized framework for fair and efficient comparisons among GFMs in the prokaryotic domain.

### 2.5 Metagenomic Annotation

For metagenomic annotation, we frame the task as a token classification problem, aiming to discern whether each token—representing a nucleotide—resides within a coding region. In this context, we fine-tuned **Gener***anno* using validated annotations from prokaryotic reference genomes. Specifically, we performed random sampling of 5% of genomic fragments from all prokaryotic genomes available in RefSeq. Importantly, this sampling was conducted at the level of individual genomic fragments rather than entire genomes, ensuring broad coverage across species. This approach offers two key advantages: (1) it enables the model to generalize across nearly all species in a few-shot manner, and (2) it mitigates the risk of overfitting or memorizing specific genomes during downstream evaluation.

To prepare the training data, for each sampled DNA fragment, we constructed two binary label sequences of equal length based on the RefSeq annotations—one corresponding to the positive strand and the other to the negative strand. Each position in these label sequences corresponds to a nucleotide in the DNA fragment. Within each sequence, a value of 1 indicates that the position is part of a coding region (CDS) on the respective strand, while 0 denotes a non-coding region. A contiguous stretch of 1s thus represents a CDS region on that particular strand. This dual-label structure is specifically designed to distinguish overlapping CDS regions located on opposite strands.

In terms of implementation, we adapted the original architecture of **Gener***anno* by replacing its Masked Language Modeling head with two parallel token classification heads, denoted as TokenCLS+ and TokenCLS-, which are responsible for predicting coding regions on the positive and negative strands, respectively. During inference, the model takes as input a single DNA strand (preferably the positive strand from 5’ to 3’, for consistency), and outputs two independent label sequences representing predicted CDS regions on both strands. The combination of these two sequences constitutes the complete annotation output. Further technical details are provided in Supplementary Section D.

#### Remark

While our dual-strand labeling scheme effectively addresses overlapping CDS regions across complementary strands, it does not resolve overlaps occurring within the same strand. For instance, if gene A spans positions 400 to 800 and gene B spans positions 600 to 1000 on the same strand, the label sequence would represent a single continuous segment from 400 to 1000, thereby preventing the model from distinguishing the two overlapping genes. Although such intra-strand overlaps are relatively rare, they represent a known biological phenomenon [39, 14] and remain an active area of our ongoing research. Nevertheless, we believe that releasing this foundational version of **Gener***anno*, despite its imperfections, provides a robust and transparent framework upon which the broader research community can build and improve collaboratively.

## 3 Experiments

### 3.1 Prokaryotic Gener Tasks

To evaluate the performance of **Gener***anno* on Prokaryotic Gener Tasks, we conducted a comprehensive comparison with several state-of-the-art genomic foundation models: DNABERT-2 [82], NT-v2 [20], GenomeOcean-500M, and GenomeOcean-4B [83]. In selecting these models, we required that their training sets include prokaryotic sequences, ensuring relevance to our benchmark. Consequently, we excluded models such as HyenaDNA [52], Caduceus [66], and **Gener***ator* [78], which are trained exclusively on eukaryotic data. While it is highly likely that **Gener***anno* would outperform these eukaryotic-focused models on Prokaryotic Gener Tasks, we believe that such comparisons would be neither fair nor meaningful, given the domain-specific nature of our benchmark. Additionally, we noted the potential relevance of HybriDNA [49], but unfortunately, we were unable to access its open-source implementation. Furthermore, we acknowledge that Evo series [51, 13] and METAGENE [46] represent valuable benchmarks for comparison. However, their inclusion in this study was precluded by the excessive computational resources required to run these models, which feature massive parameter sizes. We hope that by releasing this preliminary yet robust evaluation framework, researchers from academia and industry will engage with us to collaborate, enabling further refinement and expansion of this work.

The evaluation process consisted of two main steps: hyperparameter search and 10-fold cross-validation. During the hyperparameter search phase, we exhaustively tested all combinations of learning rates {1*e*^−5^, 2*e*^−5^, 5*e*^−5^, 1*e*^−4^, 2*e*^−4^, 5*e*^−4^} and batch sizes {64, 128, 256, 512}. The results reported in Table 2 are based on the optimal hyperparameters identified through this process, validated using 10-fold cross-validation to ensure robustness and reproducibility. For more technical details, please refer to Supplementary Section C.

**Table 2:** Performance of GFMs on Prokaryotic Gener Tasks. The reported metrics are averaged over 10-fold cross-validation, with the standard error in parentheses.

|  | DNABERT-2<br>(117M) | NT-v2<br>(500M) | GenomeOcean<br>(500M) | GenomeOcean<br>(4B) | **Generanno**<br>(500M) |
| --- | --- | --- | --- | --- | --- |
| Fitness: Minimal Media Glucose | 0.328 (0.043) | 0.838 (0.010) | 0.859 (0.007) | 0.863 (0.006) | **0.875 (0.003)** |
| Fitness: L-Histidine (nutrient) | 0.305 (0.028) | 0.815 (0.033) | 0.855 (0.006) | 0.864 (0.005) | **0.872 (0.004)** |
| Fitness: Pyruvate (C) | 0.255 (0.023) | 0.552 (0.048) | 0.734 (0.007) | 0.729 (0.034) | **0.742 (0.005)** |
| Fitness: L-Arabinose (C) | 0.385 (0.041) | 0.706 (0.015) | 0.760 (0.005) | 0.753 (0.005) | **0.770 (0.004)** |
| Fitness: Ammonium Chloride (N) | 0.277 (0.013) | 0.562 (0.017) | **0.674 (0.005)** | 0.669 (0.010) | 0.660 (0.010) |
| Fitness: D-Alanine (N) | 0.227 (0.023) | 0.473 (0.030) | 0.619 (0.016) | **0.646 (0.018)** | 0.633 (0.013) |
| Fitness: LB 10 °C | 0.270 (0.024) | 0.593 (0.033) | 0.582 (0.025) | 0.607 (0.013) | **0.613 (0.006)** |
| Fitness: LB 20 °C | 0.312 (0.022) | 0.599 (0.019) | 0.597 (0.022) | 0.617 (0.013) | **0.638 (0.007)** |
| Fitness: LB 30 °C | 0.170 (0.021) | 0.526 (0.029) | 0.560 (0.013) | **0.582 (0.014)** | 0.569 (0.012) |
| Fitness: LB pH 6 | 0.061 (0.014) | 0.477 (0.040) | 0.504 (0.040) | **0.573 (0.028)** | 0.549 (0.041) |
| Fitness: LB pH 8 | 0.035 (0.010) | 0.664 (0.031) | 0.695 (0.014) | **0.711 (0.003)** | **0.711 (0.008)** |
| Fitness: Cisplatin Stress | 0.153 (0.020) | 0.520 (0.031) | 0.563 (0.037) | 0.598 (0.044) | **0.602 (0.010)** |
| Fitness: Perchlorate Stress | 0.099 (0.032) | 0.561 (0.024) | 0.533 (0.048) | 0.588 (0.032) | **0.611 (0.013)** |
| Antibiotic Resistance | 0.904 (0.020) | 0.952 (0.006) | 0.969 (0.003) | 0.971 (0.003) | **0.972 (0.005)** |
| Bacterial Gene Classification | 0.902 (0.008) | 0.974 (0.003) | 0.964 (0.008) | 0.972 (0.004) | **0.981 (0.006)** |
| SSU Classification | 0.957 (0.007) | 0.963 (0.007) | 0.969 (0.001) | 0.970 (0.001) | **0.972 (0.001)** |
| Mixed Marker Classification | 0.632 (0.033) | 0.791 (0.007) | 0.827 (0.006) | **0.872 (0.006)** | 0.860 (0.004) |
| Random Fragment Classification | 0.649 (0.036) | 0.917 (0.004) | 0.911 (0.007) | 0.925 (0.006) | **0.941 (0.001)** |

Overall, **Gener***anno* demonstrated consistent superiority across nearly all tasks. Notably, beyond its outstanding performance at equivalent parameter scales, the most remarkable achievement of **Gener***anno* was its ability to surpass GenomeOcean-4B, a prokaryotic-focused model with over eight times its parameter size. This result underscores the efficiency and compactness of **Gener***anno*, highlighting its ability to achieve high performance without relying on excessive computational resources. One plausible explanation for the suboptimal performance of GenomeOcean-4B lies in two key aspects. First, GenomeOcean adopts the conventional all sequence training paradigm, whereas the effectiveness of the functional sequence training paradigm has been extensively validated in our prior work on **Gener***ator* [78] and further corroborated by Evo2 [13]. Second, as a causal DNA language model utilizing BPE tokenization, GenomeOcean-4B may inherit limitations similar to those observed in **Gener***ator*. Specifically, our previous experiments with **Gener***ator* revealed that BPE tokenization exhibits poor compatibility with causal DNA language modeling, potentially constraining its performance on downstream tasks.

On the other hand, DNABERT-2, a mixed-domain model trained on both eukaryotic and prokaryotic sequences, exhibited notably weaker performance. In particular, it performed almost entirely ineffectively in certain tasks, such as fitness prediction under perchlorate stress and varying pH conditions. This subpar performance can be attributed to two key factors: (1) its relatively small parameter size (117M), which constrains its ability to capture complex genomic patterns, and (2) its lack of specialization in prokaryotic DNA, as it was designed for mixed-domain applications. In contrast, NT-v2, another mixed-domain model, demonstrated commendable performance despite being less specialized for prokaryotic tasks than **Gener***anno* and GenomeOcean. Although its results were slightly inferior to those of prokaryotic-specific models, they remained competitive within the same order of magnitude.

### 3.2 Metagenomic Annotation

In this section, we evaluate the performance of **Gener***anno* in metagenomic annotation tasks by comparing it with other representative models in the field. These include traditional HMM-based approaches such as GLIMMER3 [26], GeneMarkS2 [47], MetaGeneMark2 [31], Prodigal [38], and MetaProdigal [38], as well as GFM-based methods like GeneLM [3]. Notably, another GFM-based method, SegmentNT, was excluded from the comparison because it is specifically trained on five eukaryotic species, making it unsuitable for direct comparison with models focused on prokaryotic genomes.

To ensure a fair and unbiased evaluation, we collected reference sequences from 33 prokaryotic species in RefSeq, comprising 33 chromosomes and 16 plasmids. To mitigate potential concerns about biased sample selection, these species were chosen as the union of all species tested by the methods included in this study, ensuring broad coverage and impartiality.

We evaluated six metrics to comprehensively assess model performance:

1. **Base-Pair Precision**: The proportion of correctly predicted coding nucleotides among all predicted coding nucleotides.
2. **Base-Pair Sensitivity**: The proportion of correctly predicted coding nucleotides among all true coding nucleotides.
3. **Start Accuracy**: The ratio of correctly identified CDS start positions, corresponding to transitions from 0 to 1 in the label sequences.
4. **End Accuracy**: The ratio of correctly identified CDS end positions, corresponding to transitions from 1 to 0 in the label sequences.
5. **Boundary Accuracy**: The ratio of correctly identified start and end positions for the same CDS.
6. **Exact Match Rate**: The ratio of perfectly predicted CDS regions, including both start and end positions, as well as the continuous coding region in between.

Among these metrics, **Gener***anno* demonstrated near-universal superiority, as summarized in Table 3 and 4. The only exception was base-pair precision, where it ranked second in bacterial chromosome annotation, narrowly trailing GeneLM by 0.001. However, this marginal improvement in precision was accompanied by a notable decrease in base-pair sensitivity. The HMM-based method Prodigal also achieved robust performance, outperforming all other methods except **Gener***anno*. However, it is important to note that such outstanding results are specific to single-species genomic assemblies and do not generalize to metagenomic settings, as evidenced by the substantial performance degradation observed for MetaProdigal.

**Table 3:** In-sample performance of gene annotation methods on bacterial chromosomes.

|  | Genomic Annotation |  |  | Metagenomic Annotation |  |  |  |
| --- | --- | --- | --- | --- | --- | --- | --- |
|  | Glimmer3 | GeneMarkS2 | Prodigal | MetaGeneMark2 | MetaProdigal | GeneLM | **Generanno** |
| Start Accuracy | 0.885 | 0.895 | <u>0.914</u> | 0.885 | 0.889 | 0.892 | **0.940** |
| End Accuracy | 0.884 | 0.895 | <u>0.914</u> | 0.887 | 0.889 | 0.892 | **0.940** |
| Boundary Accuracy | 0.794 | 0.815 | <u>0.848</u> | 0.797 | 0.802 | 0.816 | **0.890** |
| Exact Match Rate | 0.783 | 0.806 | <u>0.841</u> | 0.788 | 0.795 | 0.806 | **0.888** |
| Base-Pair Precision | 0.904 | <u>0.910</u> | 0.909 | 0.907 | 0.906 | **0.911** | <u>0.910</u> |
| Base-Pair Sensitivity | 0.986 | 0.989 | <u>0.992</u> | 0.991 | <u>0.992</u> | 0.943 | **0.998** |

**Table 4:** In-sample performance of gene annotation methods on bacterial plasmids.

|  | Genomic Annotation |  |  | Metagenomic Annotation |  |  |  |
| --- | --- | --- | --- | --- | --- | --- | --- |
|  | Glimmer3 | GeneMarkS2 | Prodigal | MetaGeneMark2 | MetaProdigal | GeneLM | **Generanno** |
| Start Accuracy | 0.817 | 0.840 | <u>0.846</u> | 0.834 | 0.844 | 0.829 | **0.890** |
| End Accuracy | 0.817 | <u>0.856</u> | 0.846 | 0.841 | 0.837 | 0.827 | **0.896** |
| Boundary Accuracy | 0.685 | <u>0.741</u> | 0.740 | 0.724 | 0.730 | 0.722 | **0.815** |
| Exact Match Rate | 0.673 | <u>0.733</u> | <u>0.733</u> | 0.716 | 0.720 | 0.712 | **0.813** |
| Base-Pair Precision | 0.853 | 0.873 | 0.865 | 0.870 | 0.864 | <u>0.875</u> | **0.877** |
| Base-Pair Sensitivity | 0.980 | 0.984 | <u>0.986</u> | 0.984 | 0.985 | <u>0.920</u> | **0.994** |

#### On In-Sample Performance

It is important to note that the 33 species used in this evaluation do not represent a traditional machine learning test set, i.e., data unseen during training. Instead, they reflect the in-sample model performance across multiple prokaryotic species. For instance, *Escherichia coli*, a cornerstone organism in microbiology, is widely included in the training datasets of virtually all practical annotation methods. This approach is not necessarily problematic in genomics due to the concept of phylogenetic correlation—organisms share a common evolutionary origin, and closely related species often exhibit over 95% gene similarity [43, 44]. Consequently, even if a species like *E. coli* were excluded from the training set, models could still achieve near in-sample performance by learning from its close relatives.

#### Zero-Shot Generalization Test

To rigorously evaluate the generalization capabilities of **Gener***anno* and GeneLM, we conducted a zero-shot test using archaeal genomes. Since neither model was trained on archaeal sequences, we collected 42 reference sequences from 31 archaeal species in RefSeq, comprising 31 chromosomes and 11 plasmids. As shown in Table 5 and 6, both models demonstrated robust generalization performance. However, **Gener***anno* significantly outperformed GeneLM, achieving results comparable to the in-sample performance of traditional HMM-based methods. These results indicate that **Gener***anno* is capable of generalizing to previously unseen genomic contexts, suggesting its applicability to gene annotation in yet-to-be-discovered or poorly characterized bacterial genomes. While direct validation on such targets remains to be conducted, the strong zero-shot performance on archaeal genomes provides a promising indication of its broad generalization capability.

**Table 5:** Performance of gene annotation methods on archaeal chromosomes. Models marked in red correspond to zero-shot evaluations without exposure to archaeal sequences during training.

|  | Genomic Annotation |  |  | Metagenomic Annotation |  |  |  |  |
| --- | --- | --- | --- | --- | --- | --- | --- | --- |
|  | Glimmer3 | GeneMarkS2 | Prodigal | MetaGeneMark2 | MetaProdigal | **GeneLM** | **Generanno** | **Generanno** |
| Start Accuracy | 0.856 | 0.860 | 0.870 | 0.870 | 0.860 | 0.756 | 0.874 | **0.892** |
| End Accuracy | 0.857 | 0.860 | 0.870 | 0.870 | 0.860 | 0.755 | <u>0.875</u> | **0.889** |
| Boundary Accuracy | 0.739 | 0.747 | 0.767 | 0.766 | 0.748 | 0.568 | <u>0.771</u> | **0.800** |
| Exact Match Rate | 0.730 | 0.741 | 0.762 | 0.759 | 0.741 | 0.553 | <u>0.764</u> | **0.797** |
| Base-Pair Precision | 0.846 | 0.856 | 0.853 | 0.853 | 0.851 | 0.852 | **0.858** | <u>0.855</u> |
| Base-Pair Sensitivity | 0.988 | 0.989 | <u>0.992</u> | 0.991 | 0.991 | 0.823 | 0.990 | **0.996** |

**Table 6:** Performance of gene annotation methods on archaeal plasmids. Models marked in red correspond to zero-shot evaluations without exposure to archaeal sequences during training.

|  | Genomic Annotation |  |  | Metagenomic Annotation |  |  |  |  |
| --- | --- | --- | --- | --- | --- | --- | --- | --- |
|  | Glimmer3 | GeneMarkS2 | Prodigal | MetaGeneMark2 | MetaProdigal | **GeneLM** | **Generanno** | **Generanno** |
| Start Accuracy | 0.787 | 0.802 | <u>0.844</u> | 0.832 | 0.835 | 0.723 | 0.826 | **0.868** |
| End Accuracy | 0.798 | 0.822 | <u>0.839</u> | 0.834 | 0.833 | 0.745 | 0.830 | **0.848** |
| Boundary Accuracy | 0.648 | 0.684 | <u>0.732</u> | 0.721 | 0.719 | 0.554 | 0.713 | **0.758** |
| Exact Match Rate | 0.640 | 0.678 | <u>0.728</u> | 0.712 | 0.715 | 0.549 | 0.707 | **0.749** |
| Base-Pair Precision | 0.818 | 0.841 | 0.829 | 0.839 | 0.827 | 0.834 | **0.845** | <u>0.844</u> |
| Base-Pair Sensitivity | 0.984 | 0.979 | <u>0.988</u> | 0.982 | 0.987 | 0.837 | 0.982 | **0.992** |

To further enhance its practical utility, we fine-tuned **Gener***anno* on a mixture of bacterial and archaeal genomic sequences. This formally released version (black-colored entry in Table 5 and 6) achieved improved performance on archaeal genomes. Interestingly, the zero-shot variant of **Gener***anno* exhibited slightly higher base-pair precision than the fine-tuned version. This counterintuitive observation may be attributed to our label curation strategy: during preprocessing, we excluded all hypothetical CDS regions and pseudogenes, retaining only experimentally validated coding regions. While this ensured high label accuracy, it likely led to an underestimation of base-pair precision, as many real but unvalidated coding regions were omitted. For a detailed discussion of this labeling methodology and its impact on performance estimation, please refer to Supplementary Section D.

Notably, the two subtasks from the Prokaryotic Gener Tasks—gene classification and taxonomic classification—further contribute to revolutionizing the conventional workflow of metagenomic annotation. While **Gener***anno* has already demonstrated remarkable performance in annotating coding regions, these tasks extend its capabilities to address broader challenges in metagenomics. Below, we provide a detailed discussion of their relevance and the exceptional performance of **Gener***anno* on these tasks.

#### Pseudogene Identification

Pseudogene prediction is a critical step in gene annotation, yet it remains a formidable challenge for traditional HMM-based methods. These methods struggle to capture the subtle distinctions between pseudogenes and active coding regions, often necessitating time-intensive comparative genomics and functional assays that rely on extensive reference databases [72]. To address this limitation, we assessed the performance of GFMs in predicting pseudogenes directly from raw sequence data. As shown in Table 7, all evaluated GFMs demonstrated strong capabilities in distinguishing pseudogenes from functional coding regions. More detailed confusion matrices of different GFMs on bacterial gene classification are provided in Supplementary Figure S1. Notably, **Gener***anno* achieved particularly outstanding results, underscoring its superior ability to discern fine-grained distinctions in genomic sequences. By integrating pseudogene prediction into the gene annotation pipeline, **Gener***anno* significantly simplifies the process, requiring only a single prediction per annotated gene region without the need for additional resources or complex workflows.

**Table 7:** Detailed performance of GFMs on bacterial gene classification. The reported values represent the accuracy averaged over 10-fold cross validation, with the standard error in parentheses.

|  | DNABERT-2<br>(117M) | NT-v2<br>(500M) | GenomeOcean<br>(500M) | GenomeOcean<br>(4B) | **Generanno**<br>(500M) |
| --- | --- | --- | --- | --- | --- |
| Intergenic | 0.930 (0.013) | 0.962 (0.006) | 0.963 (0.007) | 0.971 (0.008) | **0.981 (0.009)** |
| CDS | 0.868 (0.045) | 0.942 (0.006) | 0.934 (0.016) | 0.937 (0.007) | **0.947 (0.010)** |
| Pseudo | 0.627 (0.028) | 0.957 (0.009) | 0.889 (0.031) | 0.946 (0.009) | **0.962 (0.019)** |
| tRNA | 0.997 (0.001) | 0.997 (0.001) | 0.997 (0.001) | 0.997 (0.001) | **0.998 (0.002)** |
| rRNA | 0.980 (0.007) | 0.993 (0.004) | 0.992 (0.006) | 0.994 (0.003) | **1.000 (0.001)** |
| ncRNA | **0.998 (0.002)** | 0.994 (0.004) | 0.997 (0.002) | 0.997 (0.002) | **0.998 (0.003)** |

#### Taxonomic Classification

Taxonomic classification is a cornerstone of metagenomic analysis, particularly when dealing with complex samples such as sewage or soil, which contain DNA fragments from a wide variety of prokaryotic species. Traditional methods typically rely on identifying specific marker genes, such as SSU (e.g., 16S rRNA), to perform taxonomic classification. However, this process can be highly resource-intensive. For instance, in metagenomic workflows, SSU markers are often identified after assembling fragmented sequences into contigs, a step that is computationally demanding and error-prone when dealing with mixed-species datasets. In contrast, as evidenced by their performance in random fragment classification in Table 8, GFMs demonstrate the ability to directly classify species from arbitrary DNA fragments, bypassing the need for marker gene identification or genome assembly. This capability significantly streamlines the taxonomic classification process, especially in complex metagenomic datasets involving multiple species. Among the evaluated GFMs, **Gener***anno* exhibited particularly outstanding performance, showcasing its superior contextual understanding and generalization capabilities. By integrating taxonomic classification with gene annotation, **Gener***anno* offers a dual-purpose solution, enabling simultaneous species identification and gene annotation for these fragments, thereby significantly reducing the complexity of metagenomic analysis.

**Table 8:** Detailed performance of GFMs on random fragment-based taxonomic classification. The reported values represent the accuracy averaged over 10-fold cross validation, with the standard error in parentheses.

|  | DNABERT-2<br>(117M) | NT-v2<br>(500M) | GenomeOcean<br>(500M) | GenomeOcean<br>(4B) | **Generanno**<br>(500M) |
| --- | --- | --- | --- | --- | --- |
| Phylum | 0.759 (0.023) | 0.938 (0.003) | 0.938 (0.004) | 0.952 (0.005) | **0.958 (0.002)** |
| Class | 0.741 (0.022) | 0.928 (0.003) | 0.928 (0.004) | 0.944 (0.005) | **0.952 (0.002)** |
| Order | 0.576 (0.045) | 0.893 (0.004) | 0.882 (0.005) | 0.898 (0.009) | **0.918 (0.003)** |
| Family | 0.593 (0.027) | 0.897 (0.004) | 0.887 (0.006) | 0.892 (0.012) | **0.925 (0.003)** |

## 4 Discussion & Future Development

In this study, we introduced **Gener***anno*, a compact yet powerful genomic foundation model specifically optimized for metagenomic annotation. Through extensive benchmarking on the Prokaryotic Gener Tasks and comparative evaluations with state-of-the-art gene annotation methods, **Gener***anno* has demonstrated consistent superiority across nearly all tasks, establishing itself as a transformative tool in the field of prokaryotic genomics.

### 4.1 Key Contributions

First, we introduce **Gener***anno*, a compact yet highly effective genomic foundation model that excels across a wide range of prokaryotic genomic analysis tasks. Notably, despite its modest parameter size of 500M, **Gener***anno* outperforms GenomeOcean-4B [83], a model with over eight times its parameters, demonstrating exceptional performance that defies conventional scaling laws. This remarkable performance stems from our adoption of functional sequence training, a proper tokenization scheme, and meticulous architectural design that balance single-nucleotide resolution, computational efficiency, and context coverage.

In addition, we curated Prokaryotic Gener Tasks, the first benchmark suite tailored for the prokaryotic domain. Designed with three guiding principles—biological relevance, multispecies coverage, and sequence length diversity—this benchmark fills a critical gap in the field. By providing a rigorous framework for evaluating GFMs, Prokaryotic Gener Tasks facilitates fair comparisons and fosters advancements in prokaryotic genomics research.

Most importantly, **Gener***anno* revolutionizes metagenomic annotation by achieving superior performance across both traditional HMM-based methods [27, 47, 38], and other GFM-based approaches [3]. Beyond classic coding region annotations, **Gener***anno* pioneers the prediction of pseudogenes and taxonomic classification directly from raw sequence data. These capabilities significantly streamline traditional metagenomic workflows, marking a major leap forward in the integration of contextual understanding and biological utility.

### 4.2 Limitations and Future Directions

Despite its strengths, **Gener***anno* currently faces one notable limitation: it cannot effectively resolve overlapping genes located on the same strand. Although the model can identify overlapping regions, it predicts them as continuous intervals rather than separate entities, necessitating additional post-processing strategies for precise gene delineation. We regard the development of intra-strand disentanglement mechanisms as an important direction for future work. Nonetheless, we believe that releasing this foundational version of **Gener***anno* represents a valuable step toward collaborative improvement, as it provides a transparent and functional framework for the broader research community to build upon, refine, and extend across diverse biological contexts.

Looking ahead, we aim to extend **Gener***anno* to eukaryotic gene annotation, a more challenging domain due to the sparsity of coding regions and the complex exon-intron structure of eukaryotic genomes [84]. Unlike prokaryotic sequences, eukaryotic gene annotation requires not only identifying coding regions, but also determining whether multiple exons belong to the same gene. SegmentNT [24] has made pioneering efforts in this area, but it still lags behind traditional HMM-based methods like Augustus [69]. This highlights the vast potential for further optimization in eukaryotic gene annotation using GFMs.

Our broader vision is encapsulated in the Gener Project, an ongoing initiative that includes **Gener***anno* and **Gener***ator* [78]. Drawing inspiration from Mixture of Experts (MoE) architectures in LLMs [15, 45], we propose a division of expertise grounded in evolutionary relationships. MoE has emerged as a popular technique in LLMs, where tasks are dynamically allocated to specialized submodules (experts) during inference. Recently, MoE models have also begun to gain traction in the AI for science domain, as exemplified by ProGen3 [11] and xTrimoPGLM [16], which showcase their potential in analyzing complex biological data. However, unlike conventional MoE approaches, which rely on the model to automatically learn and allocate experts, our framework explicitly assigns expertise based on natural evolutionary boundaries—eukaryotic, prokaryotic, and viral domains. This deliberate design enhances interpretability by aligning model architecture with biological principles and improves computational efficiency through modular specialization.

As illustrated in Figure 3, we divide the tasks into four experts: **Gener***ator*-Eukaryote [78], **Gener***anno*-Prokaryote, and planned **Gener***ator*-Prokaryote and **Gener***anno*-Eukaryote. The **Gener***ator* experts focus on generative sequence design and long-sequence analysis (greater than 8k), while the **Gener***anno* experts specialize in gene annotation and fine-grained short-sequence analysis (less than 8k). This modular architecture allows individual models to be deployed, updated, and scaled independently, simplifying maintenance and reducing resource demands. Collectively, these four models form a ‘handcrafted MoE’, combining the strengths of different architectures to address diverse genomic challenges.

Unlike prokaryotes and eukaryotes, viral genomes lack complete biological functionality and rely on host organisms for essential life processes [18]. We argue that pre-training solely on viral sequences might be insufficient; instead, we propose a strategy of continued pre-training on host-specific models to achieve a comprehensive understanding of both viral and host sequences. This approach enables integrative analyses that capture the intricate interactions between viruses and their hosts, offering new insights into viral biology.

### 4.3 Broader Implications

The success of **Gener***anno* highlights the potential of ‘compact yet powerful’ models in AI for science. In an era where scaling laws [41] dominate the development of LLMs, the computational costs of training and deploying these models have become prohibitively high. Similarly, recent advancements in AI for science, such as ESM3 [35], ProGen3 [11], xTrimoPGLM [16], and Evo2 [13], have pushed parameter sizes into the magnitude of 100 billion, rendering these models prohibitively expensive for resource-constrained research teams. In contrast, **Gener***anno* demonstrates that task-specific optimizations and efficient architectures can deliver exceptional performance without relying on massive parameter sizes. This aligns with our belief that scientific tools should prioritize accessibility and usability, enabling private deployment and fine-tuning for specific research needs.

Recent findings from ProteinGym [53] further support this perspective, suggesting that scaling laws may not always yield meaningful improvements in scientific domains. For instance, many models achieve peak performance within the range between 500M and 10B parameters, beyond which performance plateaus or even degrades. Interestingly, this observation aligns with the parameter count estimation proposed by Sergey Ovchinnikov [79], assuming that protein language models primarily learn evolutionary couplings. One plausible explanation for this phenomenon lies in the inherent randomness of genetic mutations, which introduces background noise in biological sequences. Larger models are more prone to overfitting this noise, deviating from the functional truths encoded in genomic data [77]. This highlights a core distinction between general-purpose LLMs designed for concise human language and scientific foundation models tailored for noisy biological sequences.

Finally, the Gener Project represents a long-term commitment to advancing genomic research through collaborative innovation. We invite researchers worldwide to engage in flexible forms of collaboration, aiming to build a comprehensive suite of tools that democratize access to genomic foundation models and accelerate progress in functional genomics, metagenomics, and related fields. To this end, all materials necessary to replicate this work—including data, code, and model weights—will be made fully open-source on the GenerTeam GitHub page.

## A Supplementary Figures

**Figure S1:**
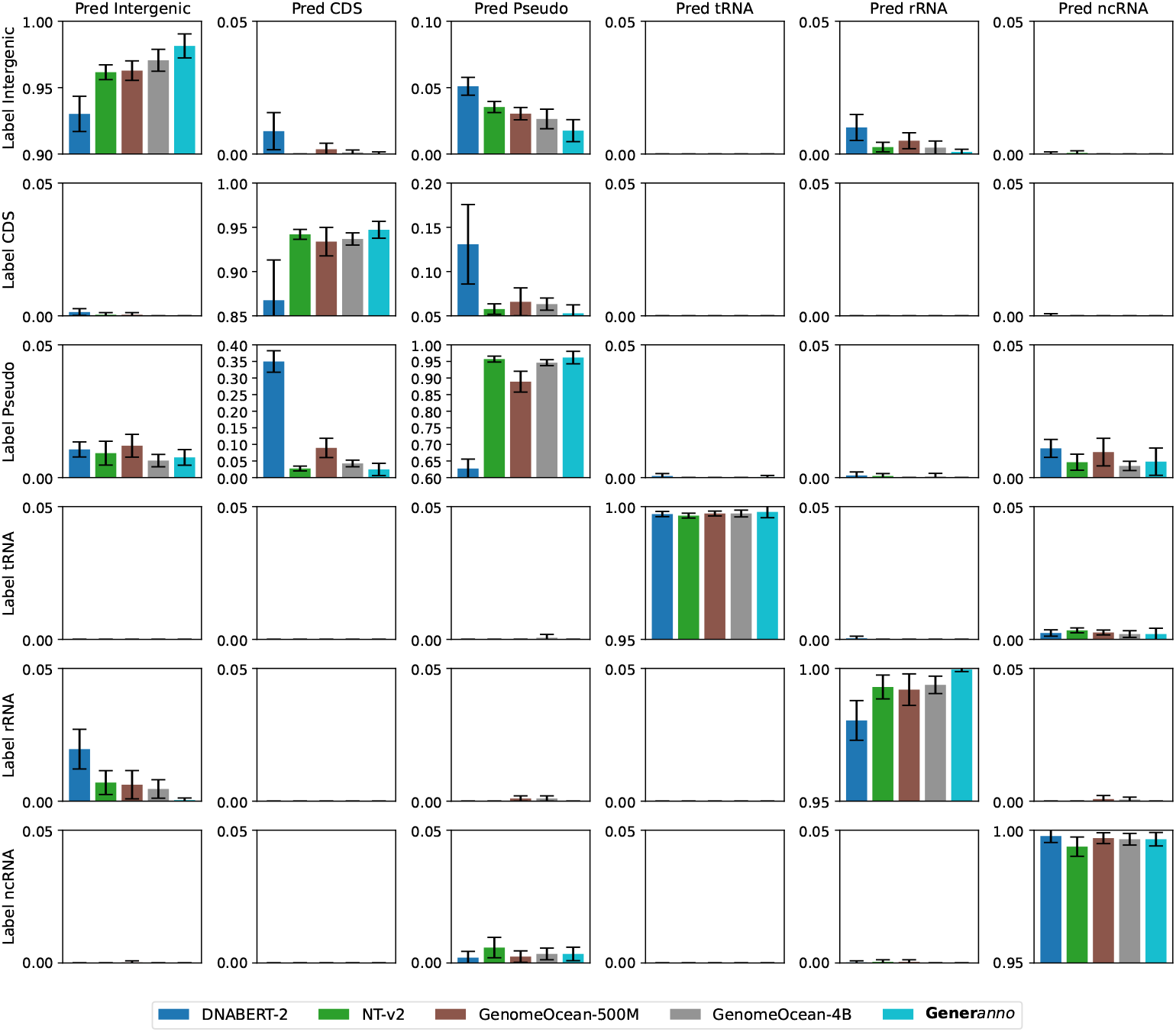
The confusion matrices of different GFMs on bacterial gene classification.

## B Details of Pre-training

In the pre-training phase, we adopted a Masked Language Modeling (MLM) objective specifically tailored for genomic sequences (`A, T, C, G, N`). Following this, 15% of the tokens in each sequence, excluding any N (which were never masked or targeted for prediction), were selected for the masking procedure. For these selected tokens, we employed a variation of the standard 8-1-1 rule [28]:

1. In 80% of cases, the selected token was replaced with a dedicated `<MASK>` token.
2. In 10% of cases, it was substituted with a randomly chosen different nucleotide from the set {`A, T, C, G`}.
3. In the remaining 10% of cases, the token was left unchanged.

An exception was made for the `<BOS>` token; if selected for masking, it was invariably replaced by the `<MASK>` token. The model was then trained to predict the original identity of these modified tokens.

For model pre-training, we employed the AdamW optimizer [48] with hyperparameters *β*_1_ = 0.9, *β*_2_ = 0.95, and a weight decay of 0.1. The learning rate schedule consisted of a linear warm-up followed by cosine decay: the learning rate increased linearly from 0 to its peak value of 4 *×* 10^−4^ over the first 2000 steps, after which it decayed according to a cosine schedule, reaching 10% of the peak value by the end of training. To ensure stable training, gradient clipping was applied with a norm threshold of 1.0. We adhered to standard practices for pre-training LLMs, using a batch size that encompassed 2 million tokens per batch. Given the maximum sequence length of 8192 tokens, this configuration resulted in batches containing 256 samples. The complete pre-training of **Gener***anno* spanned 2 epochs, with the sequence in the second epoch shifted by 4096 bp to enhance data diversity. To improve computational efficiency, we employed optimization techniques such as Flash Attention [22, 21] and the Zero Redundancy Optimizer [60, 81]. The entire pre-training process was conducted on 32 NVIDIA A100 GPUs and completed in 1040 hours.

## C Prokaryotic Gener Tasks

### C.1 Experimental Setups

To comprehensively assess the performance of **Gener***anno* across Prokaryotic Gener Tasks, we include several state-of-the-art models as baselines. For these baseline models, we conducted consistent evaluation procedures to ensure fair comparison. In our benchmark experiments, we retained the optimizer configuration from the pretraining phase, with hyperparameters *β*_1_ = 0.9, *β*_2_ = 0.95, and weight decay = 0.1. For learning rate scheduling, we adopted a ‘reduce on plateau’ strategy and implemented early stopping based on the validation dataset, with a patience of 5. The optimal learning rates and batch sizes for each model and dataset, as detailed in Table S1, were determined through an exhaustive hyperparameter search. Specifically, we evaluated all combinations of learning rates {1*e*^−5^, 2*e*^−5^, 5*e*^−5^, 1*e*^−4^, 2*e*^−4^, 5*e*^−4^} and batch sizes {64, 128, 256, 512}. For input sequences exceeding the maximum context length supported by the models, truncation was applied to fit within the constraints. For causal language models, we performed prediction through an additional linear layer using the embedding of the `<EOS>` token, while for masked language models, we used the `<BOS>` token or `<CLS>` token instead. All models obtained embeddings from their final layer and underwent full fine-tuning during the evaluation process. All evaluation metrics were obtained through 10-fold cross-validation.

These self-evaluated baseline models can be accessed through the following HuggingFace^1^ repositories:

1. DNABERT-2: `zhihan1996/DNABERT-2-117M`
2. NT-v2: `InstaDeepAI/nucleotide-transformer-v2-500m-multi-species`
3. GenomeOcean-500M: `pGenomeOcean/GenomeOcean-500M`
4. GenomeOcean-4B: `pGenomeOcean/GenomeOcean-4B`

**Table S1:**
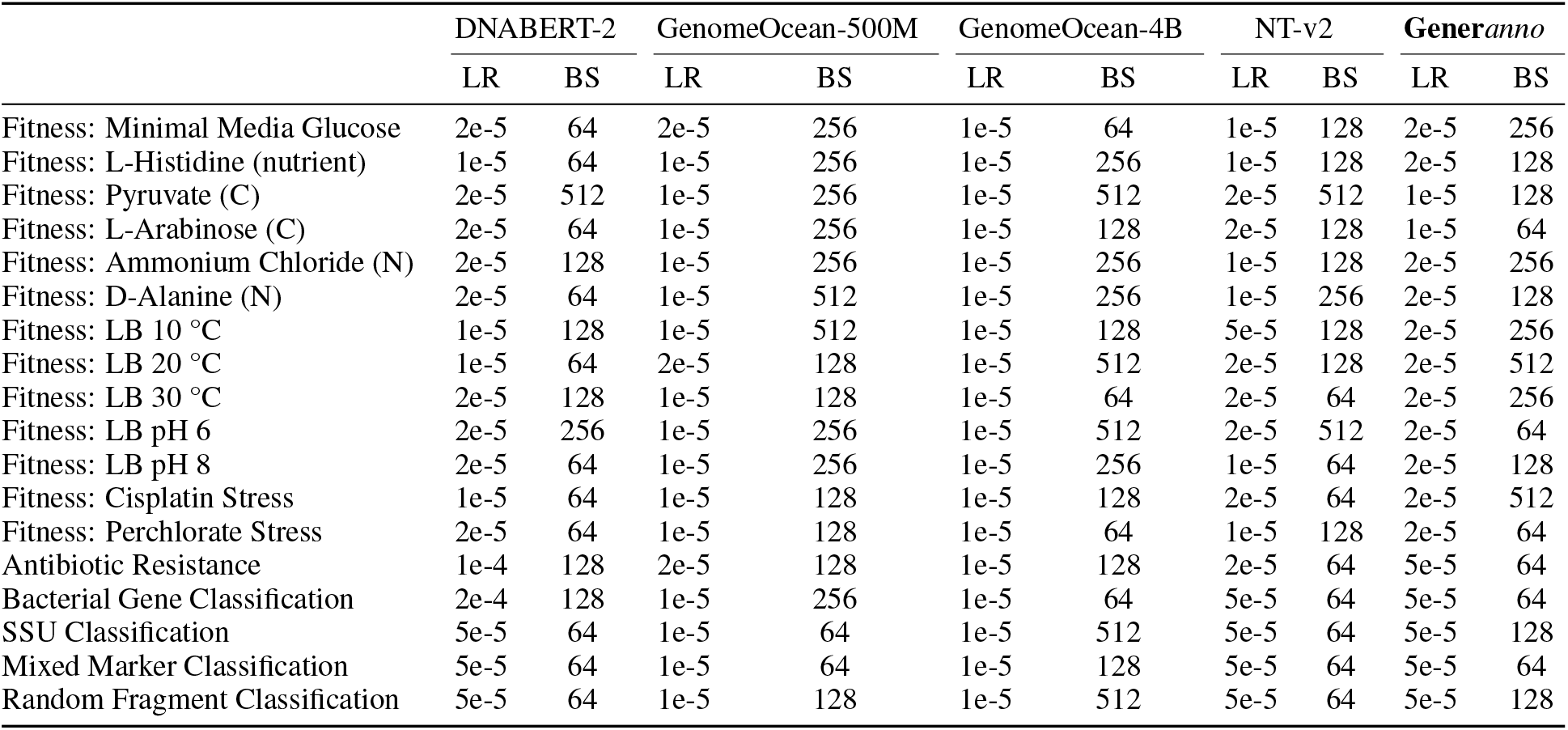
Hyperparameter settings for the Prokaryotic Gener Tasks.

|  | DNABERT-2 |  | GenomeOcean-500M |  | GenomeOcean-4B |  | NT-v2 |  | Generanno |  |
| --- | --- | --- | --- | --- | --- | --- | --- | --- | --- | --- |
|  | LR | BS | LR | BS | LR | BS | LR | BS | LR | BS |
| Fitness: Minimal Media Glucose | 2e-5 | 64 | 2e-5 | 256 | 1e-5 | 64 | 1e-5 | 128 | 2e-5 | 256 |
| Fitness: L-Histidine (nutrient) | 1e-5 | 64 | 1e-5 | 256 | 1e-5 | 256 | 1e-5 | 128 | 2e-5 | 128 |
| Fitness: Pyruvate (C) | 2e-5 | 512 | 1e-5 | 256 | 1e-5 | 512 | 2e-5 | 512 | 1e-5 | 128 |
| Fitness: L-Arabinose (C) | 2e-5 | 64 | 1e-5 | 256 | 1e-5 | 128 | 2e-5 | 128 | 1e-5 | 64 |
| Fitness: Ammonium Chloride (N) | 2e-5 | 128 | 1e-5 | 256 | 1e-5 | 256 | 1e-5 | 128 | 2e-5 | 256 |
| Fitness: D-Alanine (N) | 2e-5 | 64 | 1e-5 | 512 | 1e-5 | 256 | 1e-5 | 256 | 2e-5 | 128 |
| Fitness: LB 10 °C | 1e-5 | 128 | 1e-5 | 512 | 1e-5 | 128 | 5e-5 | 128 | 2e-5 | 256 |
| Fitness: LB 20 °C | 1e-5 | 64 | 2e-5 | 128 | 1e-5 | 512 | 2e-5 | 128 | 2e-5 | 512 |
| Fitness: LB 30 °C | 2e-5 | 128 | 1e-5 | 128 | 1e-5 | 64 | 2e-5 | 64 | 2e-5 | 256 |
| Fitness: LB pH 6 | 2e-5 | 256 | 1e-5 | 256 | 1e-5 | 512 | 2e-5 | 512 | 2e-5 | 64 |
| Fitness: LB pH 8 | 2e-5 | 64 | 1e-5 | 256 | 1e-5 | 256 | 1e-5 | 64 | 2e-5 | 128 |
| Fitness: Cisplatin Stress | 1e-5 | 64 | 1e-5 | 128 | 1e-5 | 128 | 2e-5 | 64 | 2e-5 | 512 |
| Fitness: Perchlorate Stress | 2e-5 | 64 | 1e-5 | 128 | 1e-5 | 64 | 1e-5 | 128 | 2e-5 | 64 |
| Antibiotic Resistance | 1e-4 | 128 | 2e-5 | 128 | 1e-5 | 128 | 2e-5 | 64 | 5e-5 | 64 |
| Bacterial Gene Classification | 2e-4 | 128 | 1e-5 | 256 | 1e-5 | 64 | 5e-5 | 64 | 5e-5 | 64 |
| SSU Classification | 5e-5 | 64 | 1e-5 | 64 | 1e-5 | 512 | 5e-5 | 64 | 5e-5 | 128 |
| Mixed Marker Classification | 5e-5 | 64 | 1e-5 | 64 | 1e-5 | 128 | 5e-5 | 64 | 5e-5 | 64 |
| Random Fragment Classification | 5e-5 | 64 | 1e-5 | 128 | 1e-5 | 512 | 5e-5 | 64 | 5e-5 | 128 |

### C.2 Evaluation Metrics

Regarding evaluation metrics, we adopt a unified set of metrics depending on the nature of the task. Below we summarize and define each metric.

#### C.2.1 Regression Tasks

For tasks predicting continuous values (e.g., Gene Fitness Prediction), we optimize by minimizing the **Mean Squared Error (MSE)** during training and apply early stopping based on the validation MSE. However, we report the **Pearson Correlation Coefficient (***ρ*) as a standardized measure of prediction accuracy.

The **Mean Squared Error (MSE)** measures the average squared difference between ground-truth values *y*_*k*_ and predictions *ŷ* _*k*_:

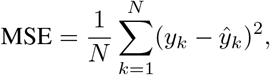

where *y*_*k*_ is the ground-truth, *ŷ* _*k*_ is the prediction for sample *k*, and *N* is the total number of samples.

The **Pearson Correlation Coefficient (***ρ*) quantifies the linear relationship between *y*_*k*_ (ground-truth) and *ŷ* _*k*_ (predicted):

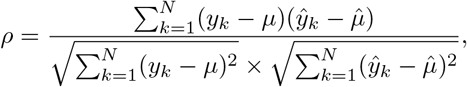

where *µ* is the mean of the ground-truth values, and 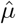 is the mean of the predicted values. While MSE ensures error minimization, *ρ* provides a complementary measure of predictive alignment, consistent with standard regression practices.

#### C.2.2 Multi-class Classification Tasks

For multi-class classification tasks (e.g., Drug Resistance Prediction, Bacterial Gene Classification), we utilize the **weighted F1 score**. This is calculated as follows:

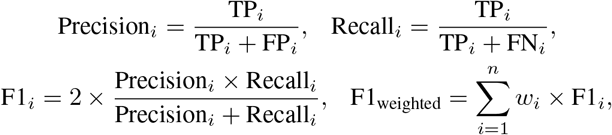

where *i* is the class index, *n* is the total number of classes, and TP_*i*_, FP_*i*_, and FN_*i*_ represent true positives, false positives, and false negatives for class *i*, respectively. The weight for class *i* is *w*_*i*_ = *n*_*i*_*/N*, where *n*_*i*_ is the number of samples in class *i*, and *N* is the total number of samples.

#### C.2.3 Multi-label Classification Tasks

For multi-label classification tasks (e.g., Taxonomic Classification), we utilize the **F1 max** metric. This metric evaluates the model performance by finding the maximum F1 score achievable across all possible thresholds for predicting labels. Specifically, the computation proceeds as follows:

1. For each label, the model outputs a prediction score indicating the likelihood of the label being positive.
2. A threshold *t* ∈ [0, 1] is applied to these scores to generate binary predictions (0 or 1).
3. For a given threshold *t*, the **F1 score** for each label is calculated as:

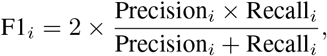

where:

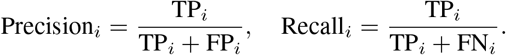
4. The **macro F1 score** is computed by averaging the F1 scores across all labels for the given threshold *t*:

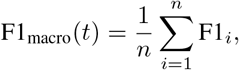

where *n* is the total number of labels, and F1_*i*_ is the F1 score for label *i* at threshold *t*.
5. Finally, the **F1 max** score is obtained by maximizing the macro F1 score across all thresholds:

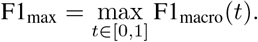

This approach ensures that the model performance is evaluated robustly, accounting for imbalances in label distribution and optimizing the threshold selection process.

## D Metagenomic Annotation

To adapt **Gener***anno* for metagenomic annotation tasks, we performed model fine-tuning, or more specifically, continued pretraining. The configuration remained consistent with the initial pretraining phase, except that the final Masked Language Model (MLM) head was replaced with the dual-strand token classification heads. Additionally, the training data was updated to better align with the gene annotation tasks, which are detailed in Section 2.5. Due to the success of the initial pretraining, the model achieved sensitivity and precision scores of 0.994 and 0.996 after the first 4000 steps during continued pretraining. Despite this impressive early performance, we continued training for a total of 88,000 steps, achieving marginal improvements. This extended training covered approximately 5% of the total nucleotide count in the RefSeq prokaryotic reference sequences.

During the evaluation phase, we downloaded 33 complete bacterial reference genomes (comprising 33 chromosomes and 16 plasmids) and 31 archaeal reference genomes (comprising 31 chromosomes and 11 plasmids). To ensure high-quality evaluation data, we filtered the reference annotation files to exclude entries labeled as hypothetical coding sequences and pseudogenes, retaining only validated CDS regions. Each annotated genomic sequence was then converted into two binary label sequences—one corresponding to the positive strand and the other to the negative strand—where each position in the label sequences corresponds to a nucleotide in the original genomic sequence. Specifically:

- A value of 1 indicates that the position is part of a validated CDS region on the respective strand.
- A value of 0 denotes either a non-coding region or an unvalidated CDS region.

This transformation enabled direct comparison with the output of the dual-strand token classification framework (Figure 2). The resulting reference annotation dataset, together with the annotation outputs of several baseline models, is publicly available at https://huggingface.co/datasets/GenerTeam/cds-annotation.

**Figure 1.**
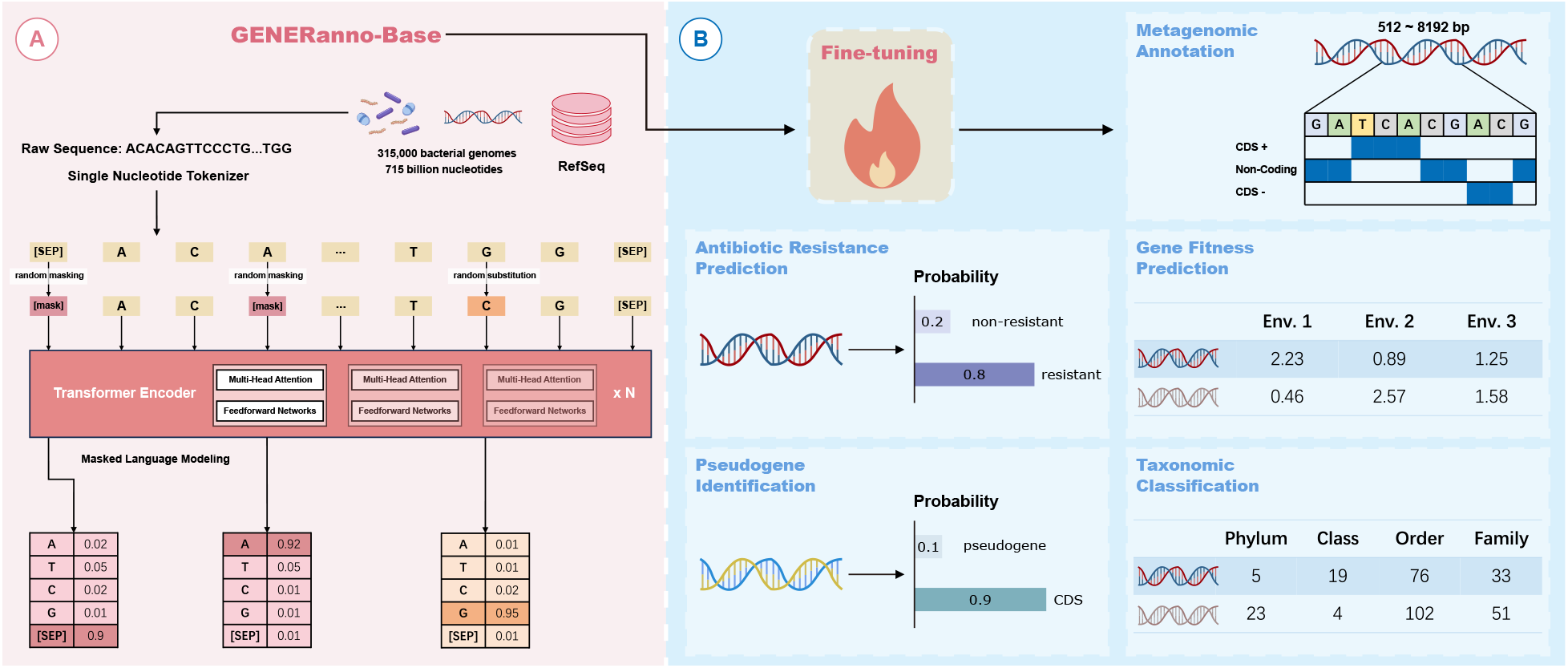
Overview of **Gener***anno*. (A) Trained on an extensive dataset comprising 715 billion nucleotides of prokaryotic DNA, **Gener***anno* employs a transformer encoder architecture with 500 million parameters, enabling bidirectional attention over sequences up to 8192 bp at single-nucleotide resolution. (B) Through fine-tuning, **Gener***anno* demonstrates its versatility across a wide range of genomic tasks, including metagenomic annotation, antibiotic resistance prediction, gene fitness prediction, pseudogene identification, and taxonomic classification.

**Figure 2.**
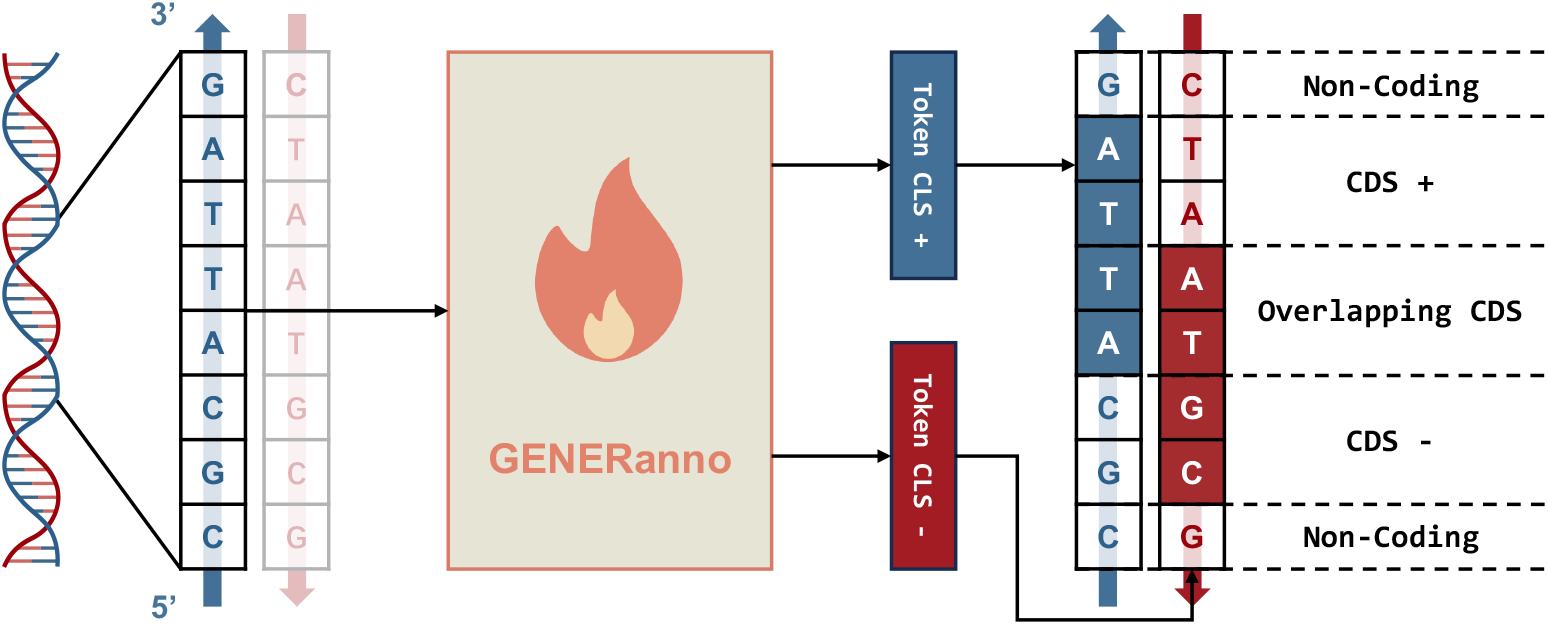
The dual-strand token classification framework for metagenomic annotation. The model takes a single DNA strand as input and predicts two independent label sequences representing coding regions on both the positive and negative strands. This structure enables the distinction of overlapping CDS regions located on opposite strands.

**Figure 3.**
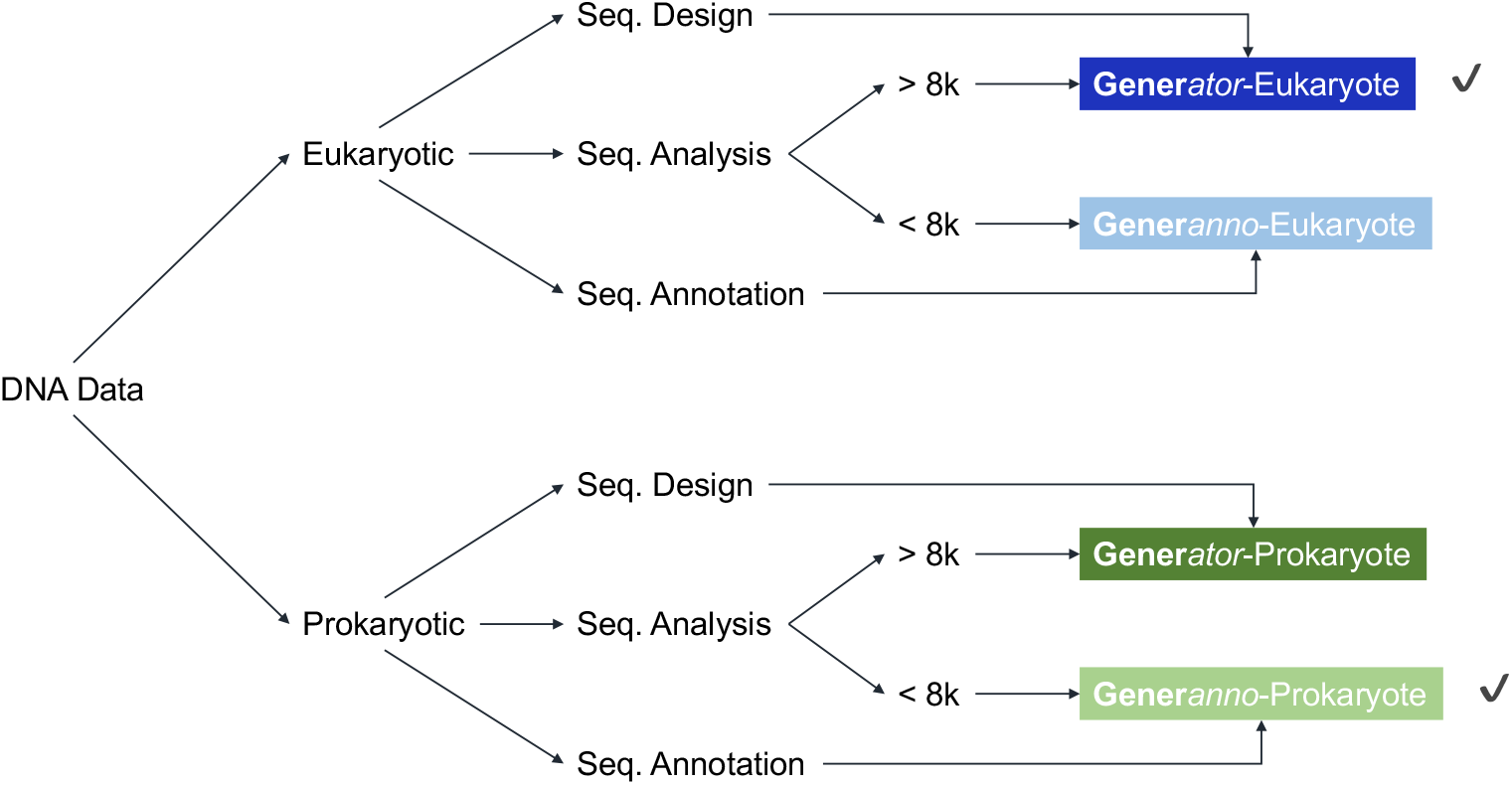
Overview of the Gener Project. This figure illustrates the modular architecture of the Gener Project, dividing tasks among four distinct experts. This design enables individual models to be deployed, updated, and scaled independently, simplifying maintenance and reducing resource demands.

The strict exclusion of hypothetical CDS regions during evaluation ensured high label accuracy but led to a lower precision score of 0.910, compared to the test precision of 0.996 achieved during continued pretraining. This discrepancy arises because hypothetical CDS regions, which are treated as non-coding regions in the evaluation, may contain a significant proportion of real but unvalidated coding regions. While this conservative filtering approach guarantees reliable and stringent evaluation metrics, it is reasonable to infer that in practical applications, the precision of **Gener***anno* could surpass the evaluated precision of 0.910.

To evaluate the performance of various annotation tools, all methods were run using their default settings. However, to accommodate the specific characteristics and requirements of each tool, different strategies were applied for processing the input sequences:

- For single-species genomic annotation tools such as GeneMarkS2, Prodigal, and Glimmer3, entire genomic sequences were provided as input to achieve optimal performance.
- For metagenomic annotation methods like MetaGeneMark2 and MetaProdigal, DNA sequences were truncated into fragments of length 8192 to simulate a metagenomic environment.
- For **Gener***anno* and GeneLM, due to differences in the maximum context lengths supported by their underlying GFMs, input sequences were automatically split into fragments of lengths 8192 and 3072 respectively.

Notably, the performance of **Gener***anno* was also tested on shorter DNA fragments to assess its robustness across varying input lengths:

- For DNA fragments of length 2048, **Gener***anno* exhibited negligible performance degradation, with sensitivity and precision dropping by less than 0.3%.
- For DNA fragments of length 512, sensitivity and precision decreased by approximately 1%.

In terms of computational efficiency, we benchmarked the runtime of these annotation methods using the reference genome of *E. coli* (approximately 4.6 million nucleotides) as an example. On a single NVIDIA RTX 4090 GPU, **Gener***anno* required approximately 1 minute to annotate the entire genome. The exact runtime varied slightly depending on the truncation length: for instance, processing with a truncation length of 8192 took around 80 seconds, while a truncation length of 2048 reduced the runtime to approximately 60 seconds. In contrast, traditional HMM-based methods such as GeneMarkS2 and Prodigal completed the task in a similar timeframe (tens of seconds), but relied on more cost-effective CPU hardware rather than GPUs.

Overall, **Gener***anno* demonstrated reliable annotation performance across DNA fragments of diverse lengths, making it a versatile tool for both genomic and metagenomic annotation tasks.

## Footnotes

1 ^1^https://huggingface.co

## References

[1] Josh Abramson, Jonas Adler, Jack Dunger, Richard Evans, Tim Green, Alexander Pritzel, Olaf Ronneberger, Lindsay Willmore, Andrew J Ballard, Joshua Bambrick, et al. Accurate structure prediction of biomolecular interactions with alphafold 3. Nature, 630:493–500, 2024.

[2] Josh Achiam, Steven Adler, Sandhini Agarwal, Lama Ahmad, Ilge Akkaya, Florencia Leoni Aleman, Diogo Almeida, Janko Altenschmidt, Sam Altman, Shyamal Anadkat, et al. Gpt-4 technical report. arXiv preprint 2303.08774, 2023.

[3] Genereux Akotenou and Achraf El Allali. Genomic language models (glms) decode bacterial genomes for improved gene prediction and translation initiation site identification. bioRxiv, 2025.

[4] Brian P. Alcock, William Huynh, Romeo Chalil, Keaton W. Smith, Amogelang R. Raphenya, Mateusz A Wlodarski, Arman Edalatmand, Aaron Petkau, Sohaib A Syed, Kara K. Tsang, Sheridan J. C. Baker, Mugdha Dave, Madeline C. McCarthy, Karyn M. Mukiri, Jalees A. Nasir, Bahar Golbon, Hamna Imtiaz, Xingjian Jiang, Komal Kaur, Megan Kwong, Zi Cheng Liang, Keyu C Niu, Prabakar Shan, Jasmine Y J Yang, Kristen L. Gray, Gemma Hoad, Baofeng Jia, Timsy Bhando, Lindsey A. Carfrae, Maya A. Farha, Shawn French, Rodion Gordzevich, Kenneth Rachwalski, Megan M. Tu, Emily Bordeleau, Damion M. Dooley, Emma J. Griffiths, Haley L. Zubyk, Eric D. Brown, Finlay Maguire, Robert G. Beiko, William W. L. Hsiao, Fiona S. L. Brinkman, Gary H. Van Domselaar, and Andrew G. McArthur. Card 2023: expanded curation, support for machine learning, and resistome prediction at the comprehensive antibiotic resistance database. Nucleic Acids Research, 51:D690–D699, 2022.

[5] Stephane Aris-Brosou and Laurent Excoffier. The impact of population expansion and mutation rate heterogeneity on dna sequence polymorphism. Molecular biology and evolution, 13(3):494–504, 1996.

[6] Pavel Avdeyev, Chenlai Shi, Yuhao Tan, Kseniia Dudnyk, and Jian Zhou. Dirichlet diffusion score model for biological sequence generation, 2023. URL https://arxiv.org/abs/2305.10699.

[7] Assaf Ben-Kish, Itamar Zimerman, Shady Abu-Hussein, Nadav Cohen, Amir Globerson, Lior Wolf, and Raja Giryes. Decimamba: Exploring the length extrapolation potential of mamba. arXiv, 2024. URL https://arxiv.org/abs/2406.14528.

[8] Johan Bengtsson-Palme, Martin Hartmann, Karl Martin Eriksson, Chandan Pal, Kaisa Thorell, Dan Göran Joakim Larsson, and Rolf Henrik Nilsson. Metaxa2: improved identification and taxonomic classification of small and large subunit rrna in metagenomic data. Molecular ecology resources, 15(6):1403–1414, 2015.

[9] John Besemer and Mark Borodovsky. Genemark: web software for gene finding in prokaryotes, eukaryotes and viruses. Nucleic acids research, 33(suppl 2):W451–W454, 2005.

[10] John Besemer, Alexandre Lomsadze, and Mark Borodovsky. Genemarks: a self-training method for prediction of gene starts in microbial genomes. implications for finding sequence motifs in regulatory regions. Nucleic acids research, 29(12):2607–2618, 2001.

[11] Aadyot Bhatnagar, Sarthak Jain, Joel Beazer, Samuel C. Curran, Alexander M. Hoffnagle, Kyle Ching, Michael Martyn, Stephen Nayfach, Jeffrey A. Ruffolo, and Ali Madani. Scaling unlocks broader generation and deeper functional understanding of proteins. bioRxiv, 2025. URL https://api.semanticscholar.org/CorpusID:277886928.

[12] Christian Bieümont and Cristina Vieira. Junk dna as an evolutionary force. Nature, 443(7111):521–524, 2006.

[13] Garyk Brixi, Matthew G Durrant, Jerome Ku, Michael Poli, Greg Brockman, Daniel Chang, Gabriel A Gonzalez, Samuel H King, David B Li, Aditi T Merchant, Mohsen Naghipourfar, Eric Nguyen, Chiara Ricci-Tam, David W Romero, Gwanggyu Sun, Ali Taghibakshi, Anton Vorontsov, Brandon Yang, Myra Deng, Liv Gorton, Nam Nguyen, Nicholas K Wang, Etowah Adams, Stephen A Baccus, Steven Dillmann, Stefano Ermon, Daniel Guo, Rajesh Ilango, Ken Janik, Amy X Lu, Reshma Mehta, Mohammad R.K. Mofrad, Madelena Y Ng, Jaspreet Pannu, Christopher Re, Jonathan C Schmok, John St. John, Jeremy Sullivan, Kevin Zhu, Greg Zynda, Daniel Balsam, Patrick Collison, Anthony B. Costa, Tina Hernandez-Boussard, Eric Ho, Ming-Yu Liu, Tom McGrath, Kimberly Powell, Dave P. Burke, Hani Goodarzi, Patrick D Hsu, and Brian Hie. Genome modeling and design across all domains of life with evo 2. bioRxiv, 2025.

[14] Gun Woo Byeon, Marc Exposit, David Baker, and Georg Seelig. Design of overlapping genes using deep generative models of protein sequences. bioRxiv, pages 2025–05, 2025.

[15] Weilin Cai, Juyong Jiang, Fan Wang, Jing Tang, Sunghun Kim, and Jiayi Huang. A survey on mixture of experts. arXiv preprint 2407.06204, 2024.

[16] Bo Chen, Xingyi Cheng, Yangli ao Geng, Shengyin Li, Xin Zeng, Bo Wang, Jing Gong, Chiming Liu, Aohan Zeng, Yuxiao Dong, Jie Tang, and Leo T. Song. xtrimopglm: Unified 100b-scale pre-trained transformer for deciphering the language of protein. bioRxiv, 2024. URL https://api.semanticscholar.org/CorpusID:259502990.

[17] Benny Chor, David Horn, Nick Goldman, Yaron Levy, and Tim Massingham. Genomic dna k-mer spectra: models and modalities. Genome biology, 10:1–10, 2009.

[18] Adrienne MS Correa, Cristina Howard-Varona, Samantha R Coy, Alison Buchan, Matthew B Sullivan, and Joshua S Weitz. Revisiting the rules of life for viruses of microorganisms. Nature Reviews Microbiology, 19(8): 501–513, 2021.

[19] Florinel-Alin Croitoru, Vlad Hondru, Radu Tudor Ionescu, and Mubarak Shah. Diffusion models in vision: A survey. IEEE Transactions on Pattern Analysis and Machine Intelligence, 45(9):10850–10869, 2023.

[20] Hugo Dalla-torre, Liam Gonzalez, Javier Mendoza Revilla, Nicolaüs Loüpez Carranza, Adam Henryk Grzywaczewski, Francesco Oteri, Christian Dallago, Evan Trop, Hassan Sirelkhatim, Guillaume Richard, Marcin J. Skwark, Karim Beguir, Marie Lopez, and Thomas Pierrot. The nucleotide transformer: Building and evaluating robust foundation models for human genomics. bioRxiv, 2024.

[21] Tri Dao. FlashAttention-2: Faster attention with better parallelism and work partitioning. In International Conference on Learning Representations (ICLR), 2024.

[22] Tri Dao, Daniel Y. Fu, Stefano Ermon, Atri Rudra, and Christopher Reü. FlashAttention: Fast and memory-efficient exact attention with IO-awareness. In Advances in Neural Information Processing Systems (NeurIPS), 2022.

[23] Bernardo P de Almeida, Franziska Reiter, Michaela Pagani, and Alexander Stark. Deepstarr predicts enhancer activity from dna sequence and enables the de novo design of synthetic enhancers. Nature genetics, 54(5):613–624, 2022.

[24] Bernardo P. de Almeida, Hugo Dalla-torre, Guillaume Richard, Christopher Blum, Lorenz Hexemer, Maxence Geülard, Javier Mendoza-Revilla, Ziqi Tang, Frederikke I. Marin, David M. Emms, Priyanka Pandey, Stefan Laurent, Marie Lopez, Alexandre Laterre, Maren Lang, Ugur Berk Sahin, Karim Beguir, and Thomas Pierrot. Annotating the genome at single-nucleotide resolution with dna foundation models. bioRxiv, 2025.

[25] Arthur L. Delcher, Douglas Harmon, Simon Kasif, Owen White, and Steven L. Salzberg. Improved microbial gene identification with glimmer. Nucleic acids research, 27 23:4636–41, 1999.

[26] Arthur L. Delcher, Douglas Harmon, Simon Kasif, Owen White, and Steven L. Salzberg. Improved microbial gene identification with glimmer. Nucleic acids research, 27 23:4636–41, 1999.

[27] Arthur L Delcher, Kirsten A Bratke, Edwin C Powers, and Steven L Salzberg. Identifying bacterial genes and endosymbiont dna with glimmer. Bioinformatics, 23(6):673–679, 2007.

[28] Jacob Devlin, Ming-Wei Chang, Kenton Lee, and Kristina Toutanova. Bert: Pre-training of deep bidirectional transformers for language understanding. In North American Chapter of the Association for Computational Linguistics, 2019.

[29] Stefan Elfwing, Eiji Uchibe, and Kenji Doya. Sigmoid-weighted linear units for neural network function approximation in reinforcement learning. Neural networks, 107:3–11, 2018.

[30] Veniamin S. Fishman, Yuri Kuratov, Maxim Petrov, Aleksei Shmelev, Denis Shepelin, N. Chekanov, Olga L. Kardymon, and Mikhail S. Burtsev. Gena-lm: a family of open-source foundational dna language models for long sequences. Nucleic Acids Research, 53, 2024.

[31] Karl Gemayel, Alexandre Lomsadze, and Mark Borodovsky. Metagenemark-2: improved gene prediction in metagenomes. BioRxiv, pages 2022–07, 2022.

[32] Katarüina Grešovaü, Vlastimil Martinek, David Čechaük, Petr Šimeček, and Panagiotis Alexiou. Genomic bench-marks: a collection of datasets for genomic sequence classification. BMC Genomic Data, 24(1):25, 2023.

[33] Albert Gu and Tri Dao. Mamba: Linear-time sequence modeling with selective state spaces. arXiv, 2024. URL https://arxiv.org/abs/2312.00752.

[34] Albert Gu, Karan Goel, and Christopher Re. Efficiently modeling long sequences with structured state spaces. In International Conference on Learning Representations, 2022.

[35] Thomas Hayes, Roshan Rao, Halil Akin, Nicholas James Sofroniew, Deniz Oktay, Zeming Lin, Robert Verkuil, Vincent Q. Tran, Jonathan Deaton, Marius Wiggert, Rohil Badkundri, Irhum Shafkat, Jun Gong, Alexander Derry, Raul S. Molina, Neil Thomas, Yousuf Khan, Chetan Mishra, Carolyn Kim, Liam J. Bartie, Matthew Nemeth, Patrick D. Hsu, Tom Sercu, Salvatore Candido, and Alexander Rives. Simulating 500 million years of evolution with a language model. bioRxiv, 2024.

[36] Yong He, Pan Fang, Yongtao Shan, Yuanfei Pan, Yanhong Wei, Yichang Chen, Yihao Chen, Yi Liu, Zhenyu Zeng, Zhan Zhou, Feng Zhu, Edward C. Holmes, Jieping Ye, Jun Li, Yuelong Shu, Mang Shi, and Zhaorong Li. Lucaone: Generalized biological foundation model with unified nucleic acid and protein language. bioRxiv, 2024.

[37] Philip Hugenholtz and Gene W Tyson. Metagenomics. Nature, 455(7212):481–483, 2008.

[38] Doug Hyatt, Gwo-Liang Chen, Philip F LoCascio, Miriam L Land, Frank W Larimer, and Loren J Hauser. Prodigal: prokaryotic gene recognition and translation initiation site identification. BMC bioinformatics, 11:1–11, 2010.

[39] Shalev Itzkovitz, Eran Hodis, and Eran Segal. Overlapping codes within protein-coding sequences. Genome research, 20(11):1582–1589, 2010.

[40] Yanrong Ji, Zhihan Zhou, Han Liu, and Ramana V. Davuluri. Dnabert: pre-trained bidirectional encoder representations from transformers model for dna-language in genome. bioRxiv, 2020.

[41] Jared Kaplan, Sam McCandlish, Tom Henighan, Tom B Brown, Benjamin Chess, Rewon Child, Scott Gray, Alec Radford, Jeffrey Wu, and Dario Amodei. Scaling laws for neural language models. arXiv preprint 2001.08361, 2020.

[42] Taku Kudo and John Richardson. Sentencepiece: A simple and language independent subword tokenizer and detokenizer for neural text processing. In Conference on Empirical Methods in Natural Language Processing, 2018.

[43] Qiuyi Li, Celine Scornavacca, Nicolas Galtier, and Yao-Ban Chan. The multilocus multispecies coalescent: A flexible new model of gene family evolution. Systematic Biology, 70(4):822–837, 11 2020. ISSN 1063-5157. doi:10.1093/sysbio/syaa084. URL https://doi.org/10.1093/sysbio/syaa084.

[44] Qiuyi Li, Yao-ban Chan, Nicolas Galtier, and Celine Scornavacca. The effect of copy number hemiplasy on gene family evolution. Systematic Biology, 73(2):355–374, 02 2024. ISSN 1063-5157. doi:10.1093/sysbio/syae007. URL https://doi.org/10.1093/sysbio/syae007.

[45] Aixin Liu, Bei Feng, Bing Xue, Bingxuan Wang, Bochao Wu, Chengda Lu, Chenggang Zhao, Chengqi Deng, Chenyu Zhang, Chong Ruan, et al. Deepseek-v3 technical report. arXiv preprint 2412.19437, 2024.

[46] Ollie Liu, Sami Jaghouar, Johannes Hagemann, Shangshang Wang, Jason Wiemels, Jeff Kaufman, and Willie Neiswanger. Metagene-1: Metagenomic foundation model for pandemic monitoring. arXiv preprint 2501.02045, 2025.

[47] Alexandre Lomsadze, Karl Gemayel, Shiyuyun Tang, and Mark Borodovsky. Modeling leaderless transcription and atypical genes results in more accurate gene prediction in prokaryotes. Genome research, 28(7):1079–1089, 2018.

[48] Ilya Loshchilov and Frank Hutter. Decoupled weight decay regularization. In International Conference on Learning Representations, 2019.

[49] Mingqian Ma, Guoqing Liu, Chuan Cao, Pan Deng, Tri Dao, Albert Gu, Peiran Jin, Zhao Yang, Yingce Xia, Renqian Luo, Pipi Hu, Zun Wang, Yuan Chen, Haiguang Liu, and Tao Qin. Hybridna: A hybrid transformer-mamba2 long-range dna language model. ArXiv, abs/2502.10807, 2025.

[50] Frederikke Isa Marin, Felix Teufel, Marc Horlacher, Dennis Madsen, Dennis Pultz, Ole Winther, and Wouter Boomsma. Bend: Benchmarking dna language models on biologically meaningful tasks. arXiv preprint 2311.12570, 2023.

[51] Eric Nguyen, Michael Poli, Matthew G. Durrant, Armin W. Thomas, Brian Kang, Jeremy Sullivan, Madelena Y Ng, Ashley Lewis, Aman Patel, Aaron Lou, Stefano Ermon, Stephen A. Baccus, Tina Hernandez-Boussard, Christopher Reü, Patrick D. Hsu, and Brian L. Hie. Sequence modeling and design from molecular to genome scale with evo. bioRxiv, 2024.

[52] Eric D Nguyen, Michael Poli, Marjan Faizi, Armin W. Thomas, Callum Birch-Sykes, Michael Wornow, Aman Patel, Clayton M. Rabideau, Stefano Massaroli, Yoshua Bengio, Stefano Ermon, Stephen A. Baccus, and Christopher Reü. Hyenadna: Long-range genomic sequence modeling at single nucleotide resolution. ArXiv, 2023.

[53] Pascal Notin. Have we hit the scaling wall for ai in science? https://pascalnotin.substack.com/p/have-we-hit-the-scaling-wall-for, 2025. Accessed: 2023-05-08.

[54] Nuala A. O’Leary, Mathew W. Wright, James Rodney Brister, Stacy Ciufo, Diana Haddad, Richard McVeigh, Bhanu Rajput, Barbara Robbertse, Brian Smith-White, et al. Reference sequence (refseq) database at ncbi: current status, taxonomic expansion, and functional annotation. Nucleic Acids Research, 44:D733–D745, 2015.

[55] Donovan H. Parks, Maria Chuvochina, Christian Rinke, Aaron J. Mussig, Pierre-Alain Chaumeil, and Philip Hugenholtz. Gtdb: an ongoing census of bacterial and archaeal diversity through a phylogenetically consistent, rank normalized and complete genome-based taxonomy. Nucleic Acids Research, 50:D785–D794, 2021.

[56] Ana Elena Peürez-Cobas, Laura Gomez-Valero, and Carmen Buchrieser. Metagenomic approaches in microbial ecology: an update on whole-genome and marker gene sequencing analyses. Microbial genomics, 6(8):e000409, 2020.

[57] Michael Poli, Jue Wang, Stefano Massaroli, Jeffrey Quesnelle, Ryan Carlow, Eric Nguyen, and Armin Thomas. Stripedhyena: Moving beyond transformers with hybrid signal processing models, 12 2023b. URL https://github.com/togethercomputer/stripedhyena, 2023.

[58] Morgan N. Price, Kelly M. Wetmore, Robert Jordan Waters, Mark Callaghan, Jayashree Ray, Hualan Liu, Jennifer V. Kuehl, Ryan A. Melnyk, Jacob S. Lamson, Yumi Suh, Hans K. Carlson, Zuelma Esquivel, Harini Sadeeshkumar, Romy Chakraborty, Grant M. Zane, Benjamin E. Rubin, Judy D. Wall, Axel Visel, Axel Visel, James Timothy Bristow, Matthew J. Blow, Adam Paul Arkin, Adam Paul Arkin, Adam M. Deutschbauer, and Adam M. Deutschbauer. Mutant phenotypes for thousands of bacterial genes of unknown function. Nature, 557: 503–509, 2018. URL https://api.semanticscholar.org/CorpusID:21708244.

[59] Lawrence R. Rabiner and Biing-Hwang Juang. An introduction to hidden markov models. IEEE ASSP Magazine, 3:4–16, 1986.

[60] Jeff Rasley, Samyam Rajbhandari, Olatunji Ruwase, and Yuxiong He. Deepspeed: System optimizations enable training deep learning models with over 100 billion parameters. In Proceedings of the 26th ACM SIGKDD International Conference on Knowledge Discovery & Data Mining, pages 3505–3506, 2020.

[61] Jason A Reuter, Damek V Spacek, and Michael P Snyder. High-throughput sequencing technologies. Molecular cell, 58(4):586–597, 2015.

[62] Subham Sekhar Sahoo, Marianne Arriola, Yair Schiff, Aaron Gokaslan, Edgar Marroquin, Justin T Chiu, Alexander Rush, and Volodymyr Kuleshov. Simple and effective masked diffusion language models, 2024. URL https://arxiv.org/abs/2406.07524.

[63] Victor Solovyevand Asaf Salamov and Asaf Solovyevand. Automatic annotation of microbial genomes and metagenomic sequences. Metagenomics and its applications in agriculture, biomedicine and environmental studies, 10:0003333703460353, 2011.

[64] Melissa Sanabria, Jonas Hirsch, Pierre M. Joubert, and Anna R. Poetsch. Dna language model grover learns sequence context in the human genome. Nat. Mac. Intell., 6:911–923, 2024.

[65] Anirban Sarkar, Ziqi Tang, Chris Zhao, and Peter K Koo. Designing dna with tunable regulatory activity using discrete diffusion. bioRxiv, 2024. doi:10.1101/2024.05.23.595630. URL https://www.biorxiv.org/content/early/2024/05/24/2024.05.23.595630.

[66] Yair Schiff, Chia-Hsiang Kao, Aaron Gokaslan, Tri Dao, Albert Gu, and Volodymyr Kuleshov. Caduceus: Bi-directional equivariant long-range dna sequence modeling. ArXiv, abs/2403.03234, 2024.

[67] Bin Shao. A long-context language model for deciphering and generating bacteriophage genomes. bioRxiv, 2024.

[68] H Ye Simon, Katherine J Siddle, Daniel J Park, and Pardis C Sabeti. Benchmarking metagenomics tools for taxonomic classification. Cell, 178(4):779–794, 2019.

[69] Mario Stanke, Rasmus Steinkamp, Stephan Waack, and Burkhard Morgenstern. Augustus: a web server for gene finding in eukaryotes. Nucleic acids research, 32(suppl 2):W309–W312, 2004.

[70] Lincoln Stein. Genome annotation: from sequence to biology. Nature reviews genetics, 2(7):493–503, 2001.

[71] Jianlin Su, Murtadha Ahmed, Yu Lu, Shengfeng Pan, Wen Bo, and Yunfeng Liu. Roformer: Enhanced transformer with rotary position embedding. Neurocomputing, 568:127063, 2024.

[72] Mitchell J Syberg-Olsen, Arkadiy I Garber, Patrick J Keeling, John P McCutcheon, and Filip Husnik. Pseudofinder: detection of pseudogenes in prokaryotic genomes. Molecular Biology and Evolution, 39(7):msac153, 2022.

[73] Hugo Touvron, Louis Martin, Kevin Stone, Peter Albert, Amjad Almahairi, Yasmine Babaei, Nikolay Bashlykov, Soumya Batra, Prajjwal Bhargava, Shruti Bhosale, et al. Llama 2: Open foundation and fine-tuned chat models. arXiv preprint 2307.09288, 2023.

[74] Evan Trop, Yair Schiff, Edgar Mariano Marroquin, Chia Hsiang Kao, Aaron Gokaslan, McKinley Polen, Mingyi Shao, Aymen Kallala, Bernardo P de Almeida, Thomas Pierrot, Yang I Li, and Volodymyr Kuleshov. The genomics long-range benchmark: Advancing DNA language models, 2025. URL https://openreview.net/forum?id=8O9HLDrmtq.

[75] Roger Waleffe, Wonmin Byeon, Duncan Riach, Brandon Norick, Vijay Korthikanti, Tri Dao, Albert Gu, Ali Hatamizadeh, Sudhakar Singh, Deepak Narayanan, Garvit Kulshreshtha, Vartika Singh, Jared Casper, Jan Kautz, Mohammad Shoeybi, and Bryan Catanzaro. An empirical study of mamba-based language models. arXiv, 2024. URL https://arxiv.org/abs/2406.07887.

[76] Benjamin Warner, Antoine Chaffin, Benjamin Clavi’e, Orion Weller, Oskar Hallström, Said Taghadouini, Alexis Gallagher, Raja Biswas, Faisal Ladhak, Tom Aarsen, Nathan Cooper, Griffin Adams, Jeremy Howard, and Iacopo Poli. Smarter, better, faster, longer: A modern bidirectional encoder for fast, memory efficient, and long context finetuning and inference. ArXiv, abs/2412.13663, 2024.

[77] Eli Weinstein, Alan Amin, Jonathan Frazer, and Debora Marks. Non-identifiability and the blessings of mis-specification in models of molecular fitness. Advances in neural information processing systems, 35:5484–5497, 2022.

[78] Wei Wu, Qiuyi Li, Mingyang Li, Kun Fu, Fuli Feng, Jieping Ye, Hui Xiong, and Zheng Wang. Generator: A long-context generative genomic foundation model, 2025. URL https://arxiv.org/abs/2502.07272.

[79] Zhidian Zhang, Hannah K Wayment-Steele, Garyk Brixi, Haobo Wang, Dorothee Kern, and Sergey Ovchinnikov. Protein language models learn evolutionary statistics of interacting sequence motifs. Proceedings of the National Academy of Sciences, 121(45):e2406285121, 2024.

[80] Wayne Xin Zhao, Kun Zhou, Junyi Li, Tianyi Tang, Xiaolei Wang, Yupeng Hou, Yingqian Min, Beichen Zhang, Junjie Zhang, Zican Dong, Yifan Du, Chen Yang, et al. A survey of large language models. ArXiv, abs/2303.18223, 2023.

[81] Yanli Zhao, Andrew Gu, Rohan Varma, Liang Luo, Chien-Chin Huang, Min Xu, Less Wright, Hamid Shojanazeri, Myle Ott, Sam Shleifer, et al. Pytorch fsdp: experiences on scaling fully sharded data parallel. arXiv preprint 2304.11277, 2023.

[82] Zhihan Zhou, Yanrong Ji, Weijian Li, Pratik Dutta, Ramana V. Davuluri, and Han Liu. Dnabert-2: Efficient foundation model and benchmark for multi-species genome. ArXiv, abs/2306.15006, 2023.

[83] Zhihan Zhou, Robert Riley, S. Kautsar, Weimin Wu, Robert Egan, Steven Hofmeyr, Shira Goldhaber-Gordon, Mutian Yu, Harrison Ho, Fengchen Liu, Feng Chen, Rachael Morgan-Kiss, Lizhen Shi, Han Liu, and Zhong Wang. Genomeocean: An efficient genome foundation model trained on large-scale metagenomic assemblies. bioRxiv, 2025. URL https://api.semanticscholar.org/CorpusID:276161724.

[84] Liucun Zhu, Ying Zhang, Wen Zhang, Sihai Yang, Jian-Qun Chen, and Dacheng Tian. Patterns of exon-intron architecture variation of genes in eukaryotic genomes. BMC genomics, 10:1–12, 2009.

